# Targeted degradation of PCNA outperforms stoichiometric inhibition to result in programed cell death

**DOI:** 10.1101/2022.01.29.478266

**Authors:** Shih Chieh Chang, Pooja Gopal, Shuhui Lim, Xiaona Wei, Arun Chandramohan, Ruban Mangadu, Jeffrey Smith, Simon Ng, Marian Gindy, Uyen Phan, Brian Henry, Anthony Partridge

## Abstract

Targeted protein degradation has emerged as a powerful technology – both as a biological tool and for broadening the therapeutic proteome. As tools to probe this approach on historically intractable targets, we have previously advanced ‘biodegraders’ — targeted degradation fusion constructs composed of mini-proteins/peptides linked to modified E3 ligase receptors. Herein, we gain deeper insights into the utility and potential of biodegraders, through a detailed study on Con1-SPOP, a biodegrader which rapidly degrades the potential cancer target, proliferating cell nuclear antigen (PCNA). In a variety of settings, the active biodegrader (Con1-SPOP) proved pharmacologically superior to its stoichiometric (non-degrading) inhibitor equivalent (Con1-SPOP_mut_). Specifically, in addition to more potent anti-proliferative effects in both 2D cell culture and 3D spheroids, PCNA degradation uniquely induced DNA damage, cell apoptosis and necrosis. Global proteomic profiling of a stable cell-line expressing Con1-SPOP under doxycycline (Dox) induction revealed that impaired mitotic division and mitochondria dysfunction is a direct consequence of PCNA degradation, effects not seen with the stoichiometric inhibitor protein. To evaluate the therapeutic potential of biodegraders, we showed that Dox-induced Con1-SPOP achieved complete tumor-growth inhibition in a xenograft model. To explore application of biodegraders as a novel therapeutic modality, modified mRNA encoding Con1-SPOP was synthesized and encapsulated into lipid nanoparticles (LNPs). The approach successfully delivered mRNA *in vitro* to deplete endogenous PCNA within hours of application and with nanomolar potency. Overall, our results demonstrate the utility of biodegraders as biological tools and highlight target-degradation as a more efficacious approach versus stoichiometric inhibition. Finally, once *in vivo* delivery and expression are optimized, biodegraders may be leveraged as an exciting therapeutic modality.

## INTRODUCTION

Targeted protein degradation (TPD) has emerged as a drug-discovery paradigm shift. The TPD approach involves the engineered recruitment of an E3 ubiquitin ligase receptor to a protein of interest (POI) to induce polyubiquitination and subsequent degradation of the POI by the 26S proteasome^1^. The related modalities based on this concept include molecular glue (e.g. Indisulam, lenalidomide)^13^, NEDDylator^4^ and small molecule based degraders^5^. Induced degradation of a POI by recruiting an E3 ligase was first demonstrated by the depletion of methionine aminopeptidase-2^6^. Protein degradation confers several advantages over stoichiometric inhibition: (1) Proteins may have several catalytic and non-catalytic functions (e.g. proteinprotein interaction/scaffolding properties) and conventional approaches typically block just one of these activities. Conversely, targeted degradation eradicates the protein and its associated functions entirely, (2) Many drug targets lack catalytic sites and are typically not addressable by traditional inhibitors; a wider range of binding sites can be leveraged as TPD is not restricted to engaging a POI at an active site, (3) Protein degradation can achieve superior pharmacological inhibition of disease relevant proteins (enzymes, transcription factors, scaffolding proteins) and achieve sustainable inhibition even after washout of the small molecule based degrader^7^, and (4) the catalytic nature of the small molecule based degrader suggests they can be effective at lower concentrations than inhibitors requiring significant target binding site saturation. This latter advantage may be particularly important for proteins with micromolar intracellular levels. Together, these advantages have generated substantial excitement. However, routine identification of well-behaved, cell-permeable, and cell-active small-molecule-based degraders has proven challenging due to their bifunctional nature, requirements for optimized linker lengths/compositions and overall large size (typically violating one or more of the Lipinski rules^8^). Indeed, the extensive medicinal chemistry required for synthesis of such molecules hamper their application as tools to address open questions in the field. For example, which of the >600 human E3 ligases are optimal for targeted degradation approaches?; and in which disease-states/tissues? Thus far, only a handful of E3 ligases have been leveraged, including Von Hippel Lindau (VHL), cereblon, cellular inhibitor of apoptosis protein 1 (cIAP1), mouse double minute 2 homolog (MDM2), DDB1-Cul4A-associated factor15 (DCAF15), DCAF16 and RNF114^9^,^10^ and X-linked inhibitor of apoptosis (XIAP) in specific and nongenetic IAP dependent protein eraser (SNIPER)^11^. In addition, application of small molecule driven approaches to classically intractable targets is hampered by a paucity of selective and high affinity POI ligands, despite the broadened number of exploitable binding sites. Accordingly, there is a need for systems that can address these gaps and advance the field. Due to its speed and versatility, the biodegrader approach offers a corresponding solution.

Here, we apply the biodegrader approach to gain insights to the advantages of TPD versus stoichiometric inhibition. Previously, we developed a platform of biodegraders^12^ – modified E3 ligases that have their substrate binding domains replaced with a high affinity small biologic (such as a nanobody, monobody, DARPin, peptide etc.) against a POI to generate a synthetic E3 ligase with engineered specificity. Similar approaches have been pursued by others under the monikers of ubiquibodies^13,14^, SCF^FNDY-B2 15^, and Ab-SPOP^16^ to degrade oncoproteins such as EGFR^17^, BCR-ABL^18^, HER2^19^, RAS^20-22^ and HIF-α^23^ using re-engineered E3 ligases such as CHIP, SPOP, FBW7, VHL, ASB1, DDB2, SOCS2 or even accessory protein HIV-1 virion infectivity factor that binds to ELOB-ELOC-CUL5^24^ Efficient degradation with this technology can be achieved using a diverse range of target binding moieties such as monobodies, nanobodies, DARPins, αREPs and peptides enabling the interrogation a wide range of protein targets, including those where identification of small molecule ligands is challenging.^12,21^. There is also flexibility in terms of the truncated E3 protein used with effective examples across several E3 classes^12,21^. Thus, the highly modular nature of biodegraders allows for rapid evaluation of substrate degradability and the corresponding functional consequences. Additional insights involve the spectrum of E3 ligases that can be leveraged, especially as they relate to subcellular localization and tissue specificity/disease-state. As a research tool, the biodegraders can be integrated into the genome with their expression controlled by a doxycycline (Dox) inducible promoter to study the consequences of targeted degradation on a time-scale that is much faster than RNAi approaches^12^. Further, the effects of protein knockdown are reversible as target levels return once the biodegrader has turned over.

An oncotarget of interest is the proliferating cell nuclear antigen (PCNA), a ubiquitously expressed protein that plays critical roles in various DNA replication processes. At the heart of the replication fork, PCNA acts as a sliding clamp and central scaffold for the dynamic engagement of replication-related proteins to influence a plethora of processes including DNA damage repair^25^ or bypass^26^, chromatin establishment^27^, sister chromatid cohesion^27^ and replication surveillance^28^. Inhibition of PCNA blocks DNA replication and results in chemo sensitization^29^ and has thus been suggested to be a broad-spectrum anti-cancer target^30^. Recently, we engineered an anti-PCNA biodegrader by replacing the substrate binding domain of the SPOP E3 ligase with a peptide called Con1 (consensus motif 1) that binds PCNA with a *K*_d_ of 100 nM^12^ and is able to inhibit DNA replication *in vitro*^31^. The resulting biodegrader, termed Con1-SPOP, builds off this activity to further induce rapid and robust degradation of PCNA and enhanced inhibition of the cell cycle and cell proliferation.

Efficient cellular delivery of biodegraders represents a major hurdle limiting their therapeutic potential. To circumvent this issue, we sought to explore LNP-mediated delivery of synthetic mRNA encoding for our PCNA biodegraders. This has been a successful strategy in delivering mRNA *in vitro*^32^ and *in vivo,* as observed with the remarkable efficacy and safety of the recent mRNA-based COVID-19 vaccines^33,34^. Recent advances in mRNA technology have made its synthesis and purification, time and cost-efficient. Compared to DNA-based therapeutic approaches, mRNA has the advantage that it poses no risk for insertional mutagenesis since it is naturally metabolized. Furthermore, mRNA therapeutics do not need to enter the nucleus, making them easier to deliver compared to DNA therapeutics.

Here, we i) explore the cellular consequences of PCNA-degradation versus PCNA-inhibition; ii) evaluate the efficacy of Con1-SPOP to inhibit tumor growth in a xenograft model; iii) determine the potency of LNP encapsulated biodegrader mRNA on endogenous PCNA in various cancer cell lines.

## RESULTS

### PCNA degradation results in DNA damage, cell apoptosis and necrosis

We previously engineered a robust anti-PCNA biodegrader Con1-SPOP (Fig. 1a), and a stoichiometric inhibitor counterpart called Con1-SPOP_mut_^12^. This latter construct deletes the three-box motif (ΔAAEILILADLHSADQLKTQAVDFIN) from SPOP to prevent formation of the ubiquitination complex. As a result, this protein stoichiometrically blocks PCNA’s function but without prompting its degradation. We also established a non-binding control, Con1_mut_-SPOP, where the Con1 sequence contains 3-point mutations (SAVLQKK<u>A</u>TD<u>AA</u>HPKK, underscored residues represent the point mutations) to abolish PCNA binding^31^. To facilitate our studies, we leveraged stable cell lines of T-REx 293 with Dox-inducible expression of either Con1-SPOP, Con1-SPOP_mut_ or Con1_mut_-SPOP^12^.

**Figure 1.**
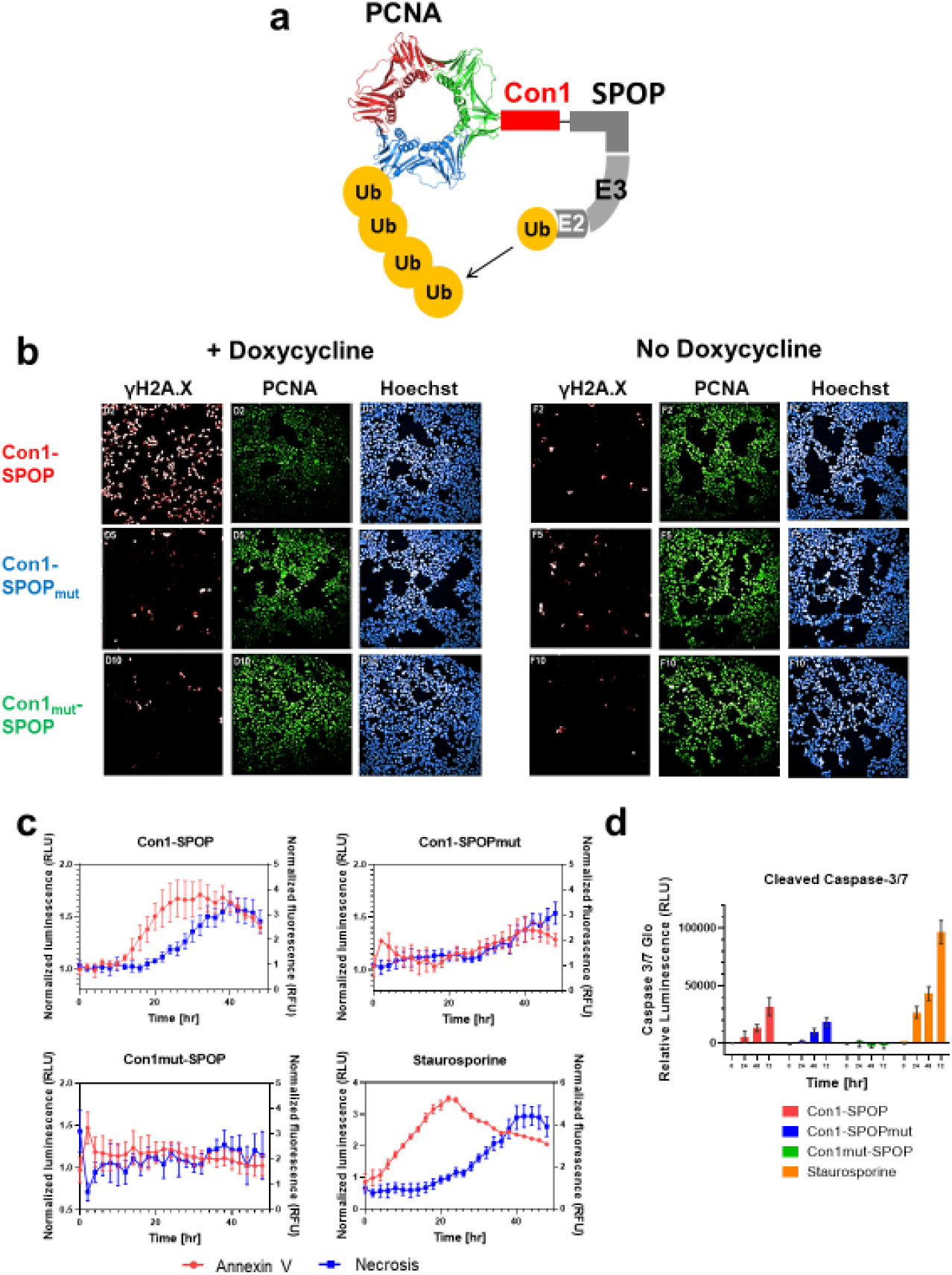
Degradation of PCNA results in DNA damage, apoptosis and necrosis. (a) Design of biodegrader Con1-SPOP for degradation of PCNA. PDB for the structures shown are PCNA (1AXC). (b) Confocal imaging of T-REx 293 cell lines stably integrated with Con1-SPOP, Con1-SPOP_mut_ and Con1_mut_-SPOP induced with 100 ng/mL of doxycycline. Immunofluorescence staining shows PCNA degradation and increased γH2A.X (Serl39) upon biodegrader expression in comparison to controls. Nuclei were counterstained with Hoechst stain. (C) Kinetic events of apoptosis and necrosis as determined by RealTime-glo™ annexin V apoptosis and necrosis assay upon induction of Con1-SPOP, Con1-SPOP_mut_ and Con1_mut_-SPOP with 10 ng/mL of doxycycline. Staurosporine at 1 μM as a positive control in T-REx 293 Con1-SPOP stable cell line. Data represents the mean of 3 replicates, normalized to the signals from no Dox control within each cell line. (d) Caspase-3/7 Glo analysis of T-REx 293 cell lines stably integrated with Con1-SPOP, Con1-SPOP_mut_ and Con1_mut_-SPOP induced with 100 ng/mL of doxycycline at different time points. Staurosporine at 1μM was used as a positive control. Data represents the mean of 3 replicates.

As PCNA plays critical roles in DNA repair, we investigated whether PCNA degradation resulted in DNA damage. Phosphorylation of histone variant H2A.X at Ser193 is a marker of DNA damage resulting from double-stranded breaks (DSBs) at sites of stalled replication forks^35,36^. Nuclear staining of γH2A.X was prominent upon Dox-induced PCNA degradation by Con1-SPOP (Fig. 1b). In contrast, these effects were not observed in cell lines expressing the stoichiometric inhibitor (Con1-SPOP_mut_) or the non-binding control (Con1_mut_-SPOP), suggesting that PCNA degradation gave superior pharmacological effects versus the stoichiometric inhibitor (Fig. 1b).

An important consideration for understanding PCNA biology, its potential as an oncology target, and the feasibility of applying the Con1-SPOP biodegrader as a therapeutic, is whether PCNA degradation/inhibition results in apoptosis and secondary necrosis. Upon Dox-induction of the biodegrader, apoptotic signals measuring Annexin V binding to phosphatidylserine (PS) on the outer leaflet of the cell membrane began to rise after 14 hr (Fig. 1c). The corresponding luminescence peaked at 32 hr and is accompanied by a rise in fluorescence signal (DNA release) indicative of membrane integrity disruption and cell death (Fig. 1c). Similar kinetics were observed upon treatment of cells with 1 μM Staurosporine, a positive control for apoptosis (Fig. 1c). In contrast, the luminescence and fluorescence signals increase concurrently after 30 hr from Dox-induced expression of PCNA inhibitor Con1-SPOP_mut_. This is not consistent with apoptotic events and likely involves alternative forms of programmed cell death. As expected, the expression of non-binding control (Con1_mut_-SPOP), did not trigger apoptosis or necrosis.

Finally, Caspase-3/7 green analysis found that the apoptotic marker continues to increase after Dox-induced expression of Con1-SPOP (Fig. 1d), 1.7 x higher than the signal detected with Con1-SPOP_mut_ with no increase in cleaved Caspase 3/7 detected for the non-binder control.

### Degradation of PCNA by Con1-SPOP leads to activation of p53

To further understand the onset of DNA damage upon PCNA depletion, we investigated additional biomarkers of genetic lesions, focusing on ataxia-telangiectasia mutated (ATM) kinase and its downstream effectors. Specifically, in response to DSB, ATM is recruited to breakage sites where it is autophosphorylated at Ser1981 and thus converting inactive ATM dimers to active monomers. ATM then phosphorylates several downstream substrates including γH2A.X (at Ser193), Check point kinase 1 (Chk1 at Ser345), Check point kinase 2 (Chk2, at Thr68) and p53 (at Ser15), modifications that regulate the cell cycle and DNA damage response^37^. In particular, in response to genotoxic stress, signaling through the ATM-Chk1/Chk2 axis transiently delays cell-cycle progression in G1/S, intra-S phase and G2/M phases^38^. To probe this pathway, we performed western blot analysis post Dox-induced (100 ng/mL) expression of Con1-SPOP and its respective controls. Consistent with the immunofluorescence results (Fig. 1b), PCNA degradation occurred as early as 4 hr and was essentially complete by 16 hr (Fig. 2a). This was followed by a time-dependent increase of γH2A.X signal. Phosphorylation of ATM (Ser1981), p53 (Ser15), Chk1 (Ser345), Chk2 (Thr68) intensified from 4 hr onwards. The same kinetics were also observed with cleaved Poly(ADP-ribose) polymerase (PARP), a marker for cells undergoing apoptosis. In contrast, we observed the downregulation of cyclin-dependent kinase inhibitor p21^Waf1/Cip1^ as PCNA degradation occurs. We also looked at proteins controlling mitotic entry/exit and cytokinesis such as Wee1, the Cdc2 inhibitory kinase, and Polo-like kinase 1 (PLK1). Wee1 kinase plays a critical role in G2/M cell cycle checkpoint arrest for the repair of damaged DNA before initiation of mitosis^39^. The mitotic kinase PLK1 is involved in processes such as mitotic entry and spindle formation^40^. Interestingly, both Wee1 and PLK1 were found to be downregulated post-Dox treatment (Fig. 2a, 2b), thus expanding the ways in which PCNA degradation can perturb the cell cycle. As for the stoichiometric control (Con1-SPOP_mut_), the Dox-induced expression resulted in a slight increase in the γH2A.X signal at 24 hr and an increase in phosphorylation of p53 (Ser15), Chk2 (Thr68), Chk1(Ser345) and downregulation of p21^Waf1/Cip1^ between 4 hr to 24 hr. While these trends were like those observed with the degrader, the intensity of the effects were weaker. Furthermore, there was no increase in cleaved PARP signal or downregulation of Wee1 or PLK1 (Fig. 2b). Dox-induced expression of the non-binder control (Con1_mut_-SPOP) showed all proteins remaining largely unperturbed, suggesting that the effects on genome integrity are specifically induced upon target engagement.

### Unbiased Global Proteomic Changes upon PCNA degradation or inhibition

As PCNA acts as a scaffolding protein for many critical functions, we sought to quantitatively assess the proteomic effects of PCNA degradation and inhibition, respectively. Three biological replicates of each stable line and the parental T-REx 293 line were conducted, with or without induction by Dox, for 4 or 24 hours, MS/MS analysis identified 4000 to 4500 unique proteins per biological replicate. A list of 3625 unique proteins were identified in all 3 replicates and used for downstream analysis (Fig. 3a). We considered a log10 (p) > 2 for proteins differentials that are statistically significant and a cut-off of log2 fold change ≥ 1 and log2 fold change ≤ 1 to indicate upregulated and downregulated proteins, respectively. As expected, in the cells expressing the active biodegrader (Con1-SPOP) after 4 hr, PCNA had the largest fold decrease in protein signal with log2 fold change of about −3 and with high statistically confidence with a −log10 (p) of 5 (Fig. 3b). On the other hand, the E3 ligase SPOP had more than a log2 fold increase of ~2 (Fig. 3b) and with high statistical significance. At 24 hr, PCNA levels continue to decrease further (Supplementary Fig. S1a).

**Figure 2.**
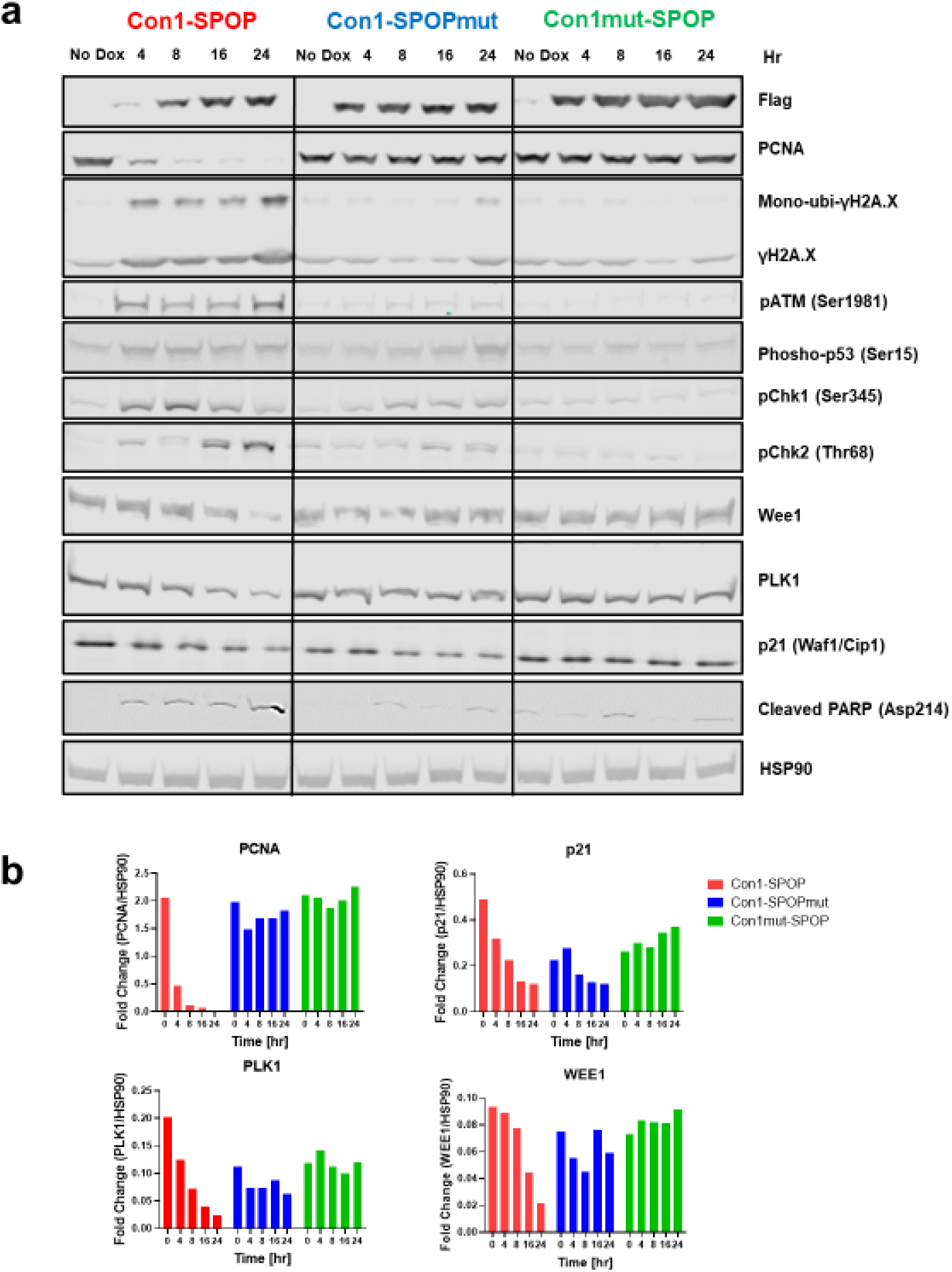
Downstream effects of targeted degradation of PCNA by biodegrader Con1-SPOP. (a) Western blot analysis of T-REx 293 cells with stable integration of Con1-SPOP (or its controls), under the control of a Tet-responsive promoter, with or without doxycycline treatment (100 ng/mL). Samples were collected at 4, 8, 16 and 24 hr. Hsp90 was used as a loading control. (b) Western band intensities of PCNA, p21^Waf1/Cip1^, Wee1 and PLK1 were quantified and normalized to the levels of the loading control Hsp90 for the respective cell lines.

Together, these results validate the approach and suggest that Dox-induced expression of Con1-SPOP rapidly degrades PCNA with high specificity. PCNA protein is unperturbed in the control and parental T-REx 293 cell lines (Supplementary Fig. S1b-g). At the 24 hr post Dox-induction time-point, PCNA levels would have been depleted for an extended period and we were interested to understand the corresponding proteomic changes. In the Con1-SPOP degrader sample, a significant upregulation of RIF1 stood out in that regard. The telomere associated protein RIF1 plays a key role in ATM/53BP1 signaling pathway and the repair of DSBs, it also promotes non-homologous end joining mediated repair and promotes replication fork protection to maintain genome stability^41^. In contrast, significant downregulation of PCNA-associated factor (PCLAF) and cell cycle proteins Kinesin-like protein KIF23, Serine/Threonine-Protein Phosphatase 2A 65 KDa Regulatory Subunit A Beta Isoform (PPP2R1B) and thymidylate synthase (TYMS) were observed (Supplementary Fig. S1a). To elucidate the temporal changes in biological patterns involved in cell cycle, DNA damage response and repair, we extracted 516 relevant proteins from the list of 3625 unique proteins base in the Reactome Knowledgebase (https://reactome.org)^42^ that are associated with cell cycle, DNA damage, DNA damage response, and apoptosis. The Z-score for the extracted list of proteins in a cell line at each time point were binned together and sorted in descending order^43^, forming six distinct clusters that are further enriched in Reactome. The clusters were rearranged to emphasize the temporal changes in each cell lines and the pathways with low false discovery rate upon enrichment were identified (Fig. 3c). From the heatmap we can observe three types of expression dynamics in Con1-SPOP and Con1-SPOP_mut_ cell line: 1) Proteins that are decreased in expression at both time points compared to no Dox control of the respective cell line, 2) Proteins that have peak expression at 4h time point, 3) Proteins that have peak expression at 24-hr time point. The first cluster was enriched for the biological pathways involving separation of sister chromatids, metaphase, anaphase and cell cycle checkpoints, suggesting that degradation or inhibition of PCNA down-regulate pathways associated with cell division at the mitotic phase. The pattern in cluster 2 suggests that at 4 hr upon PCNA degradation, there is a peak in expression of proteins involved in cell cycle checkpoint and apoptosis. The expression pattern in cluster 3 suggests that upon PCNA degradation at 24 hr, there is an upregulation of proteins related to mitochondrial translation termination, and events of apoptotic program cell death; this pattern was not observed in the stoichiometric inhibitor and other control cell lines, supporting the association of PCNA degradation and apoptosis. Upon Dox-induction at 4 hr in the PCNA stoichiometric inhibitor cell line, the biological processes were enriched for mitotic phase, chromatin organization and chromatin modifying enzymes and at 24 hr, upregulation of proteins related to transcriptional regulation by p53, mitochondrial translation processes and DNA repair was observed.

**Figure 3.**
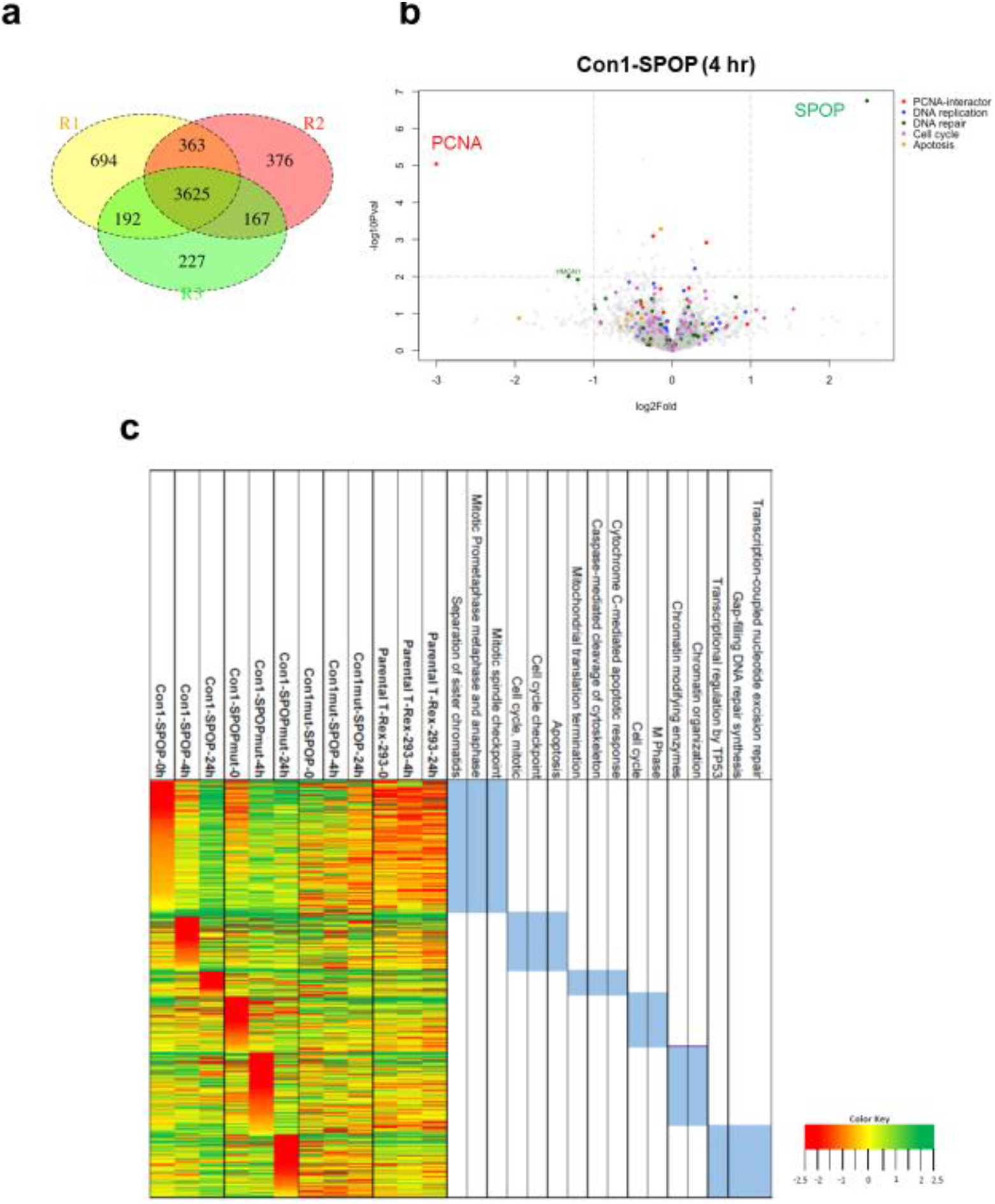
Proteomic analysis of protein differential upon PCNA degradation or inhibition. (a) Quantitative Venn diagram illustrating the overlap of common proteins identified from three biological repeats. (b) Volcano plot of protein differentials of stable cell line expressing PCNA biodegrader at 4 hr as compared to no Dox control. The x-axis presents the fold change log_2_ of and y-axis presents the p-values (-log10) of t-test between Con1-SPOP and no Dox control for individual proteins. (c) Heatmap of differentially expressed proteins. The hierarchical clustering of the 494 proteins revealed distinct patterns in six clusters. Z-score of protein abundance is color coded with red, yellow and green for high, median and low respectively (color key provided). The Reactome enriched biological pathways with the smallest false discovery rate (FDR) and functions corresponding to the relevant cluster are represented on the right. Proteomic analysis is based on 3 independent replicate experiments.

### PCNA degradation inhibits proliferation and induces cell death in three-dimensional spheroids

Next, we investigated the effect of PCNA degradation on cell proliferation. Over a four-day time period, Dox treatment (100 ng/mL) was effective in arresting the growth of the cell line expressing Con1-SPOP, slowed proliferation of the cell line expressing Con1-SPOP_mut_ but did not perturb the Con1_mut_-SPOP line (Fig. 4a). As 3D cell culture better resembles the features of *in vivo* tumors with respect to cell-cell interactions, hypoxia response and resistance^44^, we established spheroid models for stable cell lines integrated with Con1-SPOP and its respective controls. Upon Dox-induction, spheroids expressing Con1-SPOP displayed complete growth inhibition over an 80 hr time period, a proliferation pattern that resembles the 2D cell culture (Fig. 4a). Spheroids expressing Con1-SPOP_mut_ showed a slower proliferation rate upon Dox-induction, resulting in spheroids that were about 25% smaller than those without Dox-induction (Fig. 4b). Dox-induced expression of the non-binding control did not affect growth rates. Overall, the growth rates of the three Dox-induced cell lines showed a clear pattern: complete growth inhibition with the active biodegrader, marginal inhibition with the stoichiometric inhibitor, and minimal, if any growth inhibition with the non-binder control. Similar results were obtained with the CellTiter-Glo® 3D Cell Viability assay at 80 hr with cell viability inhibition measured at 72%, 18%, and 8% for the Con1-SPOP, Con1-SPOP_mut_, and Con1_mut_-SPOP constructs, respectively (Fig 4c).

### Degradation of PCNA inhibits tumor proliferation in vivo

Having established PCNA degradation as highly inhibitory towards 2D and 3D cell culture proliferation, we next evaluated the efficiency of PCNA degradation and inhibition on tumor proliferation *in vivo.* Con1-SPOP, Con1-SPOP_mut_ and Con1_mut_-SPOP stably integrated human embryonic kidney T-REx 293 cells or the parental T-REx 293 cells were injected subcutaneously into nude mice. As tumors reached an average size of 50 mm^3^, mice were assigned into 4 Dox-treated groups for efficacy and pharmacodynamic (PD) analysis, respectively (Fig. 5a). Dox-inducible degrader expression completely inhibited tumor growth in Dox-treated mice with Con1-SPOP stable cells with 7 out of 8 mice showing complete tumor regression by the end of the study at Day 30 (Supplementary Fig. S3). At Day 30, compared to the control group (Parental T-REx 293+Dox), only the active biodegrader group (Con1-SPOP + Dox) resulted in statistically significant reduction in average tumor volume, a result not seen with the stoichiometric inhibitor (Con1-SPOP_mut_ + Dox) or the non-binder groups (Con1_mut_-SPOP) (Fig. 5b). None of the groups showed significant body weight changes over the duration of the study (Fig. 5c). While the western blot analysis of tumors harvested 3 days after Dox induction showed no significant changes in PCNA levels for the Con1-SPOP + Dox group, these tumors expressed significantly higher Cleaved Caspase-3 levels as compared to the control groups (Fig. 5d and Fig. 5e). This suggests that apoptotic cell death is rapidly induced upon expression of Con1-SPOP, which could be contributing to overall decrease in protein turnover (lower GAPDH and FLAG, confounding the PCNA levels) for the Con1-SPOP + Dox group (Fig. 5d). Taken together, PCNA degradation by Con1-SPOP results in a rapid and complete inhibition of tumor growth *in vivo* as compared to stoichiometric inhibition by Con1-SPOP_mut_ or the non-binder control Con1_mut_-SPOP.

**Figure 4.**
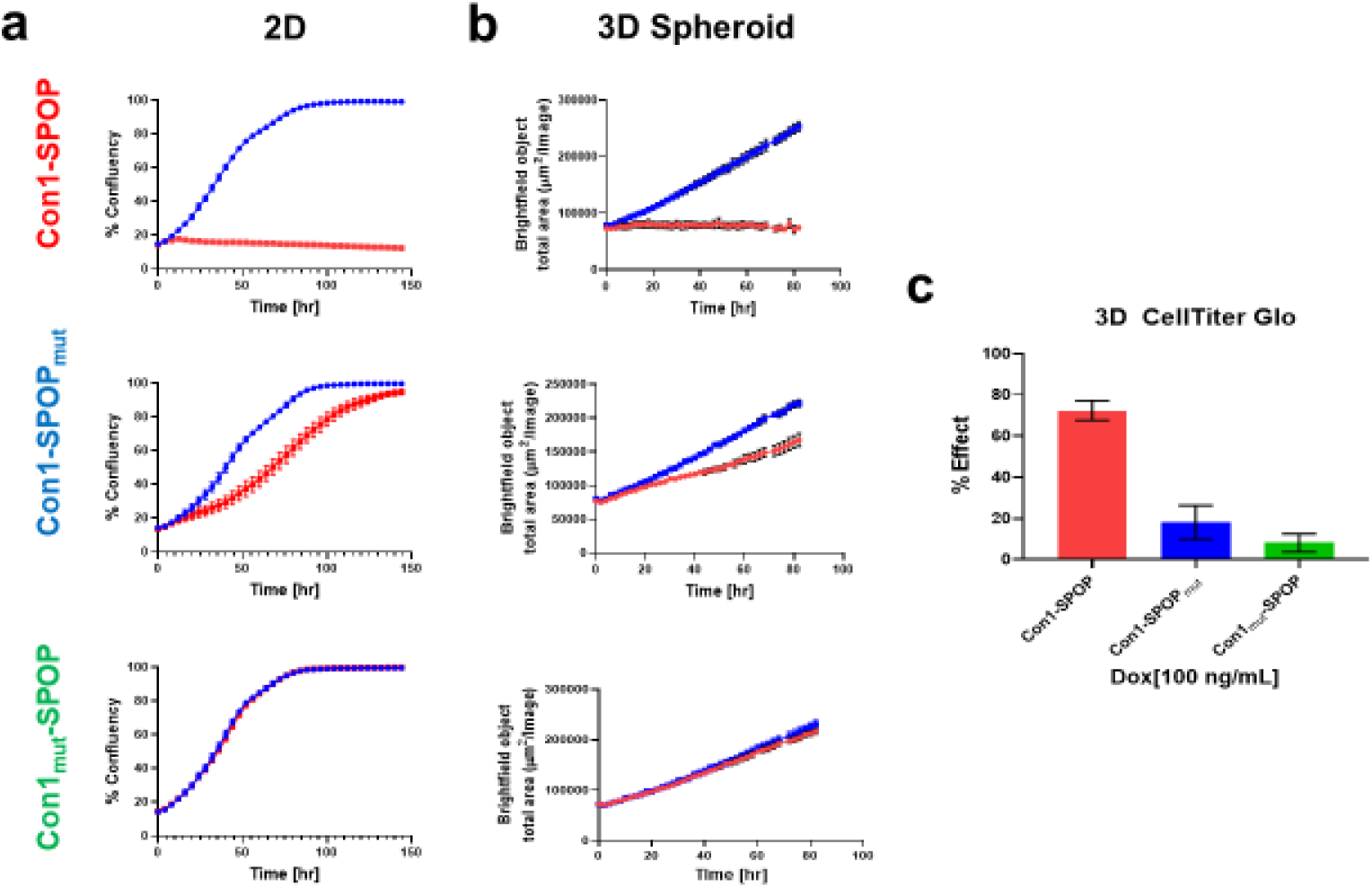
Effect of PCNA biodegrader on cell proliferation. Assessment of Dox-inducible Con1-SPOP and its control on 2D-adherent and 3D spheroids proliferation (a) Incucyte confluency measurement of 2Dadherent T-REx 293 cells with stable integration of Con1-SPOP (or its controls) with (red circles) or without (blue squares) 100 ng/mL doxycycline. (b) Spheroid proliferation curves for T-REx 293 cells with stable integration of Con1-SPOP (or its controls) with (red circles) or without (blue squares) 100 ng/mL doxycycline as determined by Incucyte. (c) Effects of PCNA degradation and inhibition in 3D spheroids on cell viability quantified by CellTiter-Glo 3D. Percent effect is determined with respect to no Dox control for respective cell lines. All experiments were performed in triplicates.

**Figure 5.**
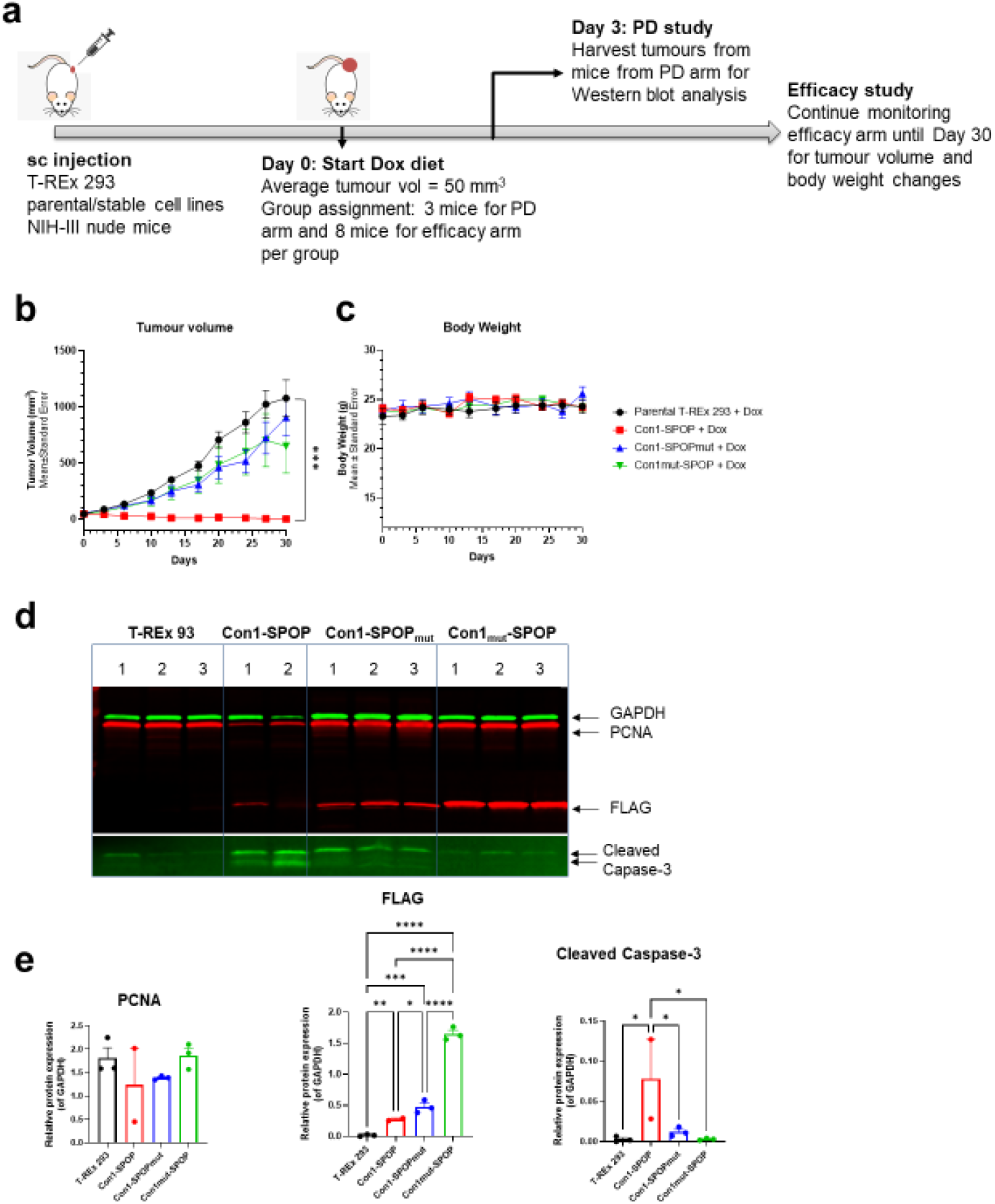
Anti-tumour effects of Con1-SPOP in the T-REx 293 xenograft model. (a) Scheme of tumor inoculation [subcutaneous (s.c.)] and treatment schedule in parental or stable cell line T-REx 293 tumourbearing NIH-III nude mice. After tumors reached an average of 50 mm^3^, mice were randomized and divided into Efficacy and PD arms: Dox-treated groups for Con1-SPOP, Con1-SPOP_mut_, Con1_mut_-SPOP and parental T-REx 293 respectively. Each efficacy and PD arm consists of *n* = 8 *and n = 3* mice respectively. (b) Individual tumor growth kinetics for efficacy arm for respective groups. Data is shown as mean tumour volume ± SEM and significance was determined by one-way ANOVA multiple comparison for tumour volumes at day 30 (*** *P* < 0.001). (c) Mean bodyweight of mice for efficacy arm from respective groups (*n* = 8). (d) Western blot analysis of PCNA, FLAG-Tag, Cleaved Caspase-3 and GAPDH (loading control) for tumor lysates from resected tumors of the PD arms harvested at day 3. For the Con1-SPOP group, 1/3 tumours in the PD arm had completely regressed by day 3 (e) Western band intensities for PCNA, FLAGTag and Cleaved Caspase-3 for the different groups normalized to levels of loading control GAPDH in respective groups. Significance was determined by ordinary one-way ANOVA multiple comparisons. *p<0.05, **p<0.01, ***p<0.001, ****p<0.0001.

### Transfection of in vitro-transcribed mRNA encoding for the biodegrader led to rapid PCNA degradation

As an intracellular biologic, cytoplasmic delivery is anticipated to be the most challenging hurdle for achieving *in vivo* activity with biodegraders. To overcome these challenges, a corresponding mRNA sequence could be delivered to the site of interest. Examples with *in vivo* activity include vaccine approaches^45^, including against Covid-19^34,46^, secreted antibodies^47^, enzyme replacement therapy^48^ and p53 replacement therapy^14,49^. Indeed, some precedence already exists for delivery of biodegrader equivalents into the mouse ear^14^. Successful cytosolic delivery of mRNA would allow the encoding degraders to achieve TPD through the ubiquitin-proteasome pathway (Fig. 6a). As an initial evaluation of this strategy, we used commercial transfection reagents to introduce mRNA of GFP degrader vhhGFP4-SPOP_167-374_ into a Dox-inducible H2B-GFP stable cell line to quantify the degradation of a model substrate, H2B-GFP. Transfection of GFP degrader mRNA reduced the number of GFP^+^ cells from 96% to basal levels (0.4%) within 24 hr (Supplementary Fig. 2a). Measuring the GFP fluorescence intensity over time showed a dramatic reduction in GFP, with a protein half-life of about 1.24 hr (Supplementary Fig. 2b). Western blot confirmed that H2B-GFP was almost undetectable by 24 hr and remained depleted at 48 hr (Supplementary Fig. 2c). Next, we synthesized mRNA encoding Con1-SPOP and the Con1-SPOP_mut_ and Con1_mut_-SPOP controls (Fig. 6b). Upon transfection in HEK293 cells with Con1-SPOP mRNA, PCNA was completely degraded by 24 hr, as assessed by western blot (Fig. 6b). PCNA degradation also resulted in reduced Histone H3 phosphorylation, indicating reduced proliferation (Fig. 6b). Indeed, transfection of anti-PCNA degrader mRNA led to strong proliferation inhibition of cancer lines (A375, A427 and AsPC-1) and immortal cell lines such as HEK293 (Fig. 6c). It is important to note that the control mRNA (Con1_mut_-SPOP) also had some effects on cell proliferation (most evident in AsPC-1) and began to recover after 48 hr, attributable to activation of the innate immune system by dsRNA contaminants likely present in the mRNA preparation. Biodegrader-mediated PCNA degradation was also compared to knockdown by siRNA. With 25nM of siRNA in A375 cells, PCNA turnover is gradual and more apparent at 96 hr (Fig. 6d), in agreement with the relatively long PCNA protein half-life of about 78.5 hr^50^. Interestingly, neither PCNA knockdown by siRNA nor the siRNA controls caused activation of Caspase-3 or cleaved PARP (Fig. 6d).

**Figure 6.**
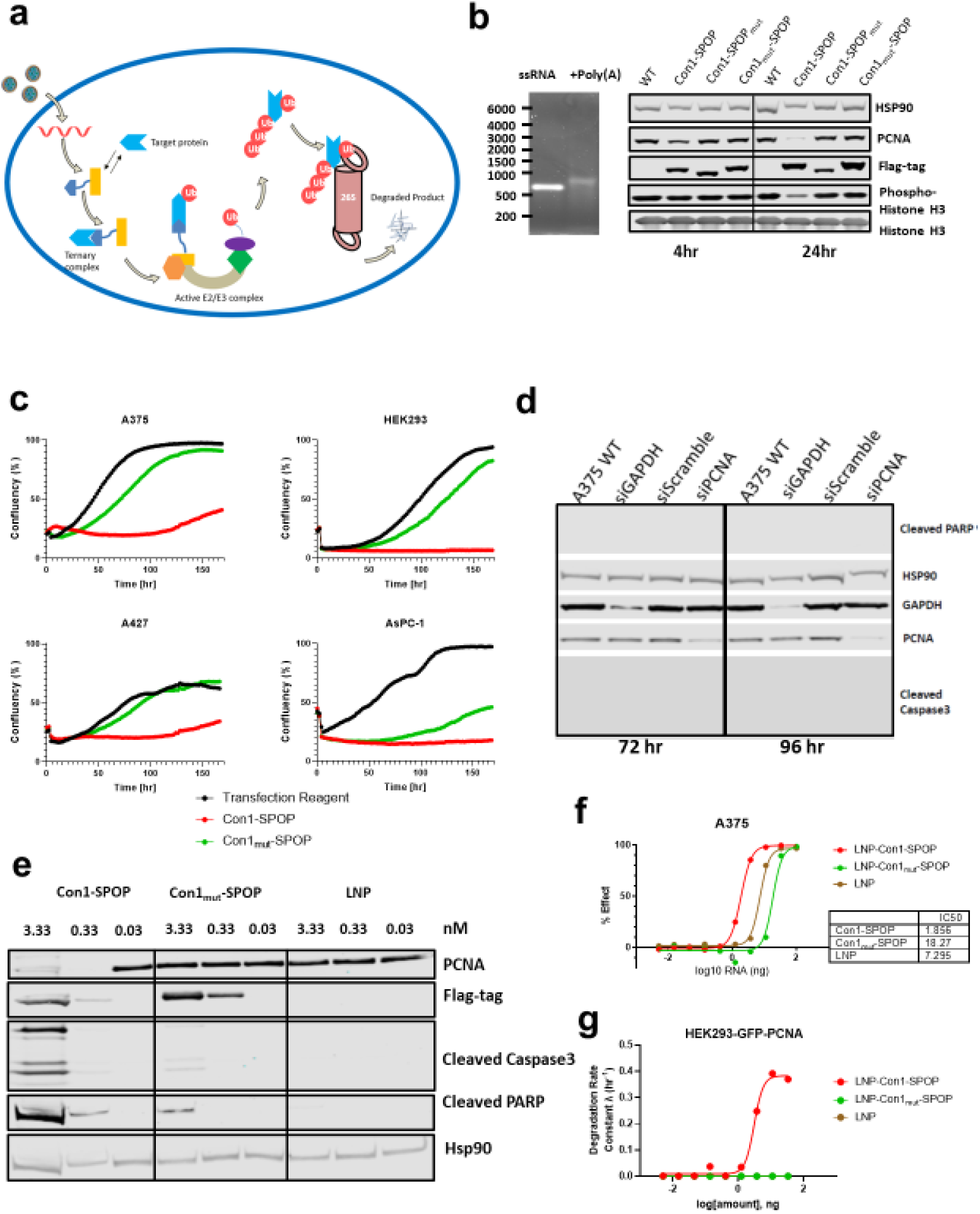
PCNA degradation via *in vitro* transcribed biodegrader mRNA. (a) Schematic of biodegrader mRNA delivery through lipid nanoparticles. (b) 1% RNA agarose gel of Con1-SPOP mRNA before and after polyadenylation (left). Western blot analysis of endogenous PCNA degradation in HEK293 cells with Con1-SPOP and its control mRNAs at 100ng per well (right). (c) Proliferation curves of AsPC-1, HEK293, A427 and A375 cells transfected with biodegrader mRNA and the non-binder control at 50 ng per well. The percentage confluency of the cells was tracked continuously over 7 days using Incucyte. (d) Western blot analysis of A375 cells treated with siRNA for PCNA knockdown. siGAPDH and siScramble at 25 nM final concentration were used as positive and negative controls respectively. (e) Western blot analysis of PCNA degradation with LNP encapsulated degrader and non-binder control mRNAs in A375 cells. Empty LNP treated cells were used as a negative control and Hsp90 as a loading control (f) Cell viability determination using CellTiterGlo viability assay in A375 cells treated with a dose titration of empty LNPs or LNPs loaded with biodegrader or non-binding control mRNA (g) Degradation kinetics of LNP-degrader and its control were determined by continuously monitoring green fluorescence over 8 hr in GFP-PCNA expressing HEK293 cells using Incucyte. Degradation rate constant was determined by fitting to a one-phase decay model using GraphPad Prism.

### Delivery of degrader mRNA using lipid nanoparticles led to rapid PCNA degradation

It has been previously shown that lipid nanoparticles can be used to encapsulate mRNA and act as a delivery vehicle^51,52^. LNPs comprise of 1) ionizable cationic lipids which promotes self-assembly for nucleic acid encapsulation and endosomal escape, 2) polyethylene glycol-lipid (PEG-lipid) that provides steric stabilization, impacts adsorption of proteins and membrane interactions, 3) Cholesterol and phospholipids that modulate physical properties such as rigidity, stability and membrane interactions. Upon encapsulation, these components protect the mRNA payload from omnipresent ribonucleases. Once taken into the cell by endocytosis, the protonation of the LNP pH-responsive lipids promotes endosomal escape of the mRNA cargo. Treatment of A375 cells with 3.3 nM of LNP-encapsulated Con1-SPOP mRNA resulted in a near complete PCNA degradation at 24 hr and induction of classical apoptotic effectors, cleaved PARP and cleaved Caspase-3 (Fig. 6e). At 0.33 nM of Con1-SPOP mRNA, we observed rapid PCNA degradation as well but with a weaker apoptotic response. Treatment with Con1_mut_-SPOP mRNA at 3.33 nM resulted in no change in PCNA levels but induced some weak apoptotic effector induction, possibly due to non-specific mRNA/LNP toxicity observed at higher concentrations. Cell viability effects of A375 treated with LNP-Con1-SPOP mRNA were assessed with CellTiter-Glo 2.0 assay and a 10-fold difference was observed in IC50 values of the active biodegrader and the non-binding control (Fig. 6f). However, empty LNPs at high concentration induce cell toxicity (Fig. 6f), which was also observed in other cancer cell lines (Supplementary Fig. S4). To further quantify the degradation kinetics of the PCNA biodegrader, LNP-Con1-SPOP mRNA was added to GFP-PCNA expressing HEK-293 cells and the GFP fluorescence was monitored over an 8 hr period. The degradation kinetic profile showed a rapid and dose-dependent degradation of GFP-PCNA (Fig. 6g).

## DISCUSSION

Traditionally, small molecule inhibitors have been successful at targeting enzymes and receptors with distinct hydrophobic binding pockets. However, targets addressable by traditional small molecules represent only about 10 to 15% of the human proteome. The remaining fraction is comprised of scaffolding proteins, PPI partners, and transcription factors that represent valuable targets that have so far resisted therapeutic intervention. Targeted protein degradation by small molecule approaches expand the scope of proteins that can be addressed since the molecular entity can be effective through binding sites that do not necessarily need to be critical for the POIs biological activity. However, small molecule-based degraders have important limitations since significant medicinal chemistry resources are required for their discovery and targets that are challenging to ligand with small molecules could be out of reach. Biodegraders offer an alternative protein silencing strategy that overcomes the limitations associated with small molecule approaches. A central advantage of biodegraders is their relatively facile discovery as they can be constructed with small-biologic POI binders that can either pre-exist or be identified with a variety of methods such as yeast- and phage-display technologies. Indeed, this has allowed us to engineer a variety of active biodegraders to GFP-fusion proteins^12^, RAS^20,22^, and PCNA^12^.

In this study, we investigated the advantages of biodegrader-mediated PCNA degradation over stoichiometric inhibition. The degradation of PCNA induced DNA damage was evident by the presence of γH2A.X, triggering phosphorylation of ATM/Chk1/Chk2, activation of p53 pathway that subsequently led to apoptosis and secondary necrosis. Meanwhile, the oncotargets Wee1 and PLK1 were found to be downregulated. This could be useful in targeting cancers with aberrant G1 checkpoint as down-regulation of Wee1 and PLK1 with resultant dysregulated G2 checkpoint, leads to mitotic lethality. In addition to its cell cycle and DNA repair functions, Chk2 also has a pro-apoptotic function, mediated in part by p53^53–55^. This is supported in part by global proteomic profiling where the protein differentials showed that PCNA degradation induced mitochondrial dysfunction and caspase-mediated events at the 24 hr time point. In terms of repair processes upon PCNA degradation, RIF1 upregulation was observed in proteomic analysis. During DNA damage, the protein RIF1 translocates to damage sites through ATM associated 53BP1 phosphorylation^56^, and suppresses chromosomal instability by stabilizing DSB ends^41^. In addition, RIF1 also facilitates BLM chromatin association and localization at DSB sites to promote DNA end processing^56^. In contrast, PCNA inhibition by the stoichiometric inhibitor Con1-SPOP_mut_ led to activation of Chk1/2-p53 but not γH2A.X and ATM. PCNA inhibition did not affect Wee1 or PLK1 protein level. As observed from proteomic profiling, PCNA inhibition results in upregulation of chromatin modifying enzymes at 4 hr and proteins related to transcription-coupled nucleotide excision repair at 24 hr. For the stoichiometric inhibitor, the kinetics of apoptotic and necrosis events upon PCNA inhibition occurred concurrently (Fig 1C), suggesting differences in the anti-proliferative cellular mechanisms as compared to PCNA degradation.

The potency of biodegrader-mediated degradation over PCNA inhibition becomes apparent when the effect on cell proliferation is compared. In both 2D and 3D cell cultures, degradation of PCNA completely stops cell proliferation and significantly reduces cell viability. Stoichiometric inhibition of PCNA slowed the proliferation rate but was unable to achieve complete inhibition. These results highlight the differences in anti-proliferative mechanisms and temporal onset of cell death. This phenomenon was directly translated to our xenograft mouse model based on Dox-inducible stable cell lines. Expression of Con1-SPOP completely inhibited tumor progression (Fig. 5b) and complete regression in 7/8 mice (Supplementary Fig. S3). In contrast, established Con1-SPOP_mut_ tumors appeared to be resistant to PCNA inhibition. The superior efficacy in the of PCNA degradation versus stoichiometric inhibition in 2D, 3D, and *in vivo* settings may be related to several phenomenon. First, PCNA has a variety of binding partners and may have several distinct biological activities. Binding of the Con1 peptide may only inhibit a subset of these. On the other hand, degradation would ablate all PCNA activities and may explain the superior efficacy. As well, degradation could be effective at sub-stoichiometric amounts of the degrader whereas the Con1-SPOP_mut_ may be limited if it cannot fully saturate and inhibit the PCNA target. In addition, the rapid kinetics of PCNA degradation may prevent the cell from adopting compensatory mechanisms related to PCNA inhibition. Alternatively, a subpopulation of tumor cells with less copies of Con1-SPOP_mut_ gene developed resistance against the inhibition, outgrowing the other population of cells. Overall, our result suggests degradation of PCNA represents a viable approach to inhibit tumor growth *in vivo* and highlights the anti-PCNA biodegrader as a potential cancer therapeutic, if efficient tumor-targeted delivery can be achieved.

To realize the therapeutic potential of biodegraders, the intracellular delivery challenge will need to be overcome. One strategy to circumvent this hurdle is to use mRNA to deliver the gene product. mRNA is efficacious in both dividing and non-dividing cells and is independent of cell cycle status. The rapid and transient expression of protein from mRNA makes it more predictable than plasmid DNA or viral vectors. Here, we synthesized anti-GFP biodegrader mRNA and showed that upon transfection, H2B-GFP gets rapidly degraded with a half-life of 1.24 hr. Next, we synthesized anti-PCNA biodegrader mRNA and showed it depletes most PCNA by 24 hr and completely inhibits proliferation in A375 and A427 cells upon mRNA transfection. A direct comparison was made with PCNA knockdown by siRNA and found that the slow turnover of PCNA did not induce apoptotic markers, suggesting protein knockdown strategies that rely on natural protein turnover rate may not be as efficacious as rapid targeted degradation. Among potential delivery vehicles, LNP encapsulation of mRNA shows promise with the recent advances in the vaccine space^57–59^. The LNPs can be synthesized in a scalable manner while shielding mRNA against degradation by ribonucleases. Furthermore, they can potentially be directed to a desired cell type using surface decoration with cell-specific ligands^60^ and subsequently facilitate mRNA endosomal escape. One example of successful therapeutic application of LNP mediated RNA delivery is Onpattro, a siRNA therapeutic for amyloidogenic transthyretin (TTR) amyloidosis that has recently gained FDA approval^61^. We encapsulated biodegrader and non-binder mRNAs within LNPs. Upon treatment in A375 cells, we observed robust degradation of PCNA with 3.33 and 0.33 nM of the anti-PCNA biodegrader and induction of apoptosis marker cleaved Caspase-3 within 24 hr. We noted that at high mRNA concentrations, the control mRNA and LNP alone can induce cell toxicity as reflected by a cell viability assay. While these results collectively demonstrate the superiority of PCNA degradation, there remains a need to further optimize the mRNA construct and delivery strategies to obtain sufficient delivery to the tumor in order for *in vivo* therapeutic effects to be realized.

Strategies for enhancing molecular stabilization include engineering the mRNA sequence to increase secondary structure in the coding sequence and 3’-UTR^62^, to boost translation rates and protein expression levels. One can also consider incorporating sets of 5’ and 3’-UTRs to increase protein expression^63^. A key element for mRNA optimization is avoiding activation of various innate immunity pathways such as pattern-recognition receptors (PRRs), pathogen-associated molecular patterns (PAMPs), protein kinase R (PKR), retinoic acid-inducible gene I (RIG-I) and interferon-induced tetratricopeptide repeat (IFIT). Single stranded mRNA can activate Toll-like receptors (TLRs) 7 and 8, and double stranded RNA (dsRNA) can be sensed by PKR, TLR3, 17 and 18^64^. The activation of TLRs can increase inflammation and inhibit protein translation^65,66^. It was shown that high performance liquid chromatography (HPLC) can reduce immunogenicity of the mRNA and improve translation, reduce inflammatory response as it remove contaminants such as dsRNA^67,68^. In addition, uridine depletion and incorporation of N1-methyl pseudouridine modifications have been shown to be less immunogenic and leads to improved protein translation^68,69^. Self-amplifying mRNA which encodes proteins enabling RNA replication, it encodes RNA genome of a single-stranded RNA virus such as alphavirus, a flavivirus^70^. They can be used to increase duration and level of expression. Furthermore, off-target toxicity of potent biodegraders can be prevented by incorporation of self-destruct mi-RNA target sites in the 3’-UTR, which would permit expression only in the target tissue type of interest such as hepatocellular carcinoma cells but not healthy hepatocytes^71^.

This study is a proof-of-concept demonstrating the superiority of target degradation over target inhibition employing biodegraders targeting PCNA both *in vitro* and *in vivo*. We also demonstrate how the biodegrader approach could be used as biological tools to answer questions relating to downstream effects upon degrading or inhibiting a specific target protein-of-interest. Finally, we share the therapeutic potential of this approach using LNP-encapsulated mRNA with activity in a few well-characterized cancer cell lines. Using biodegraders to target and destroy disease-associated proteins may unlock the remaining undruggable proteome and could be exploited to treat previously intractable diseases.

## MATERIALS AND METHODS

### Plasmids

The generation of plasmids for FLAG-vhhGFP4-SPOP_167-374_, FLAG-vhhGFP4mut-SPOP_167-374_, FLAG-vhhGFP4-SPOP_mut_, FLAG-Con1-SPOP, FLAG-Con1-SPOP_mut_, FLAG-Con1_mut_-SPOP, H2B-GFP, PCNA-GFP were detailed in an earlier publication^12^. For IVT of mRNA, FLAG-vhhGFP4-SPOP_167-374_, FLAG-vhhGFP4mut-SPOP_167-374_, FLAG-vhhGFP4-SPOP_mut_, FLAG-Con1-SPOP, FLAG-Con1-SPOP_mut_, FLAG-Con1_mut_-SPOP were subcloned into pEF6 (Thermo Fisher Scientific) using BamHI and NotI such that the CDS is downstream of T7 promoter. To generate doxycycline-inducible plasmids, H2B-GFP, FLAG-Con1-SPOP, FLAG-Con1-SPOP_mut_, FLAG-Con1_mut_-SPOP were subcloned into pcDNA™4/TO (Thermo Fisher Scientific) as described previously^12^. All plasmids were verified by sequencing at 1st BASE.

### Cell culture and transfection

Human embryonic kidney T-REx 293 cells which express the tetracycline repressor protein were purchased from Thermo Fisher Scientific and cultured in Eagle’s Minimum Essential Medium (MEM) GlutaMAX™ supplemented with 10% Tet system approved FBS (Clontech) and 5 μg/mL blasticidin. Stable cell lines expressing H2B-GFP, Con1-SPOP, Con1-SPOP_mut_ and Con1_mut_-SPOP have been described previously^12^. Briefly, T-REx 293 cells were selected using 400 μg/mL Zeocin 3 days post-transfection and maintained in 200 μg/mL Zeocin once stable colonies are formed. A375, A427, AsPC-1, SNU-387, HEK293 and PLC/PRF/5 cells were purchased from ATCC. A-375 cells were cultured in Dulbecco’s modified Eagle’s Medium (Gibco). AsPC-1 and SNU-387 cells were cultured in RPMI-1640 medium (Gibco). A-427, HEK293 and PLC/PRF/5 cells were cultured in MEM GlutaMAX™ (Gibco). All media were supplemented with 10% Fetal Bovine Serum (HyClone). All cells were maintained at 37°C, 5% CO_2_ and 90% relative humidity.

### Apoptosis and cell viability assays

Cells were seeded in 96-well poly-D-lysine coated μCLEAR® plates (Greiner) and allowed to attach overnight. Staurosporine at 1μM was added to T-REx 293 cells as positive control. 10 or 100 ng/mL of doxycycline were added to the cells to induce expression of anti-PCNA biodegrader and its controls. RealTime-glo™ annexin V apoptosis and necrosis reagents (Promega) were diluted with cell culture media 10% Fetal Bovine Serum (FBS) to twice the concentration of bioluminescent annexin reagents, 1 mM CaCl2 and necrosis detection reagent and added directly into the well and bioluminescent and fluorescence signals were measured by Spark 10M (Tecan). The plates were maintained at 37°C, 5% CO_2_ and 90% relative humidity. For the Caspase 3/7 Glo (Promega) assay, at the indicated time points, equal volumes of the assay reagent were added as per manufacturer’s instruction and incubated for 30 min at room temperature with shaking. For CellTiter-Glo (Promega) assay, the assay reagents were added at the designated time point as per manufacturer’s instruction followed by 10 min incubation at room temperature. Luminescence was recorded using the Tecan M1000 plate reader.

### Label-free proteomics sample preparation and analysis

Stable cell lines Con1-SPOP, Con1-SPOP_mut_, Con1_mut_-SPOP and parental T-REx 293 cells were seeded on 6-well plates and allowed to incubate overnight. The media was removed and replaced with fresh media containing 100 ng/mL Dox or no Dox. Upon Dox-induction at 4 hr and 24 hr, cells were trypsinized and washed with cold PBS twice and centrifuged to collect the cell pellet. 250 μL of pre-chilled 8M urea, Halt™ protease inhibitor cocktail (ThermoFisher) and phosphatase inhibitors (Roche) to each sample of 1-2 million cells. The cell lysates were pulse sonicated on ice 5 times with 2.5 amplitude with a cycle 5s on and 5s off. Samples were centrifuged for 10 min at 16,000 rpm. The protein concentration of the lysate was quantified using bicinchoninic acid (BCA) assay (ThermoFisher). 100 μg of protein from each sample were transferred and final concentration of 20 mM ammonium bicarbonate, 5 mM of DTT and 20 mM of iodoacetamide were added. Samples were digested with LysC for 3 hr before adding trypsin for overnight digestion at 37°C, followed by inactivation with 100 % TFA. The digested peptides were desalted with Oasis μElution plate following manufacturer’s protocol and dried using Evaporex. The peptides were solubilized with 0.1% formic acid followed by injection into LC/MS for all three biological replicates. The proteins identified in all three biological replicates were used for downstream analysis. For each of the cell lines, we identified the differentially expressed protein at each time points by comparing to the no Dox control in each cell line to generate volcano plots. p-values are calculated with t-test calculated between two technical replicates base on abundance of each protein detected by LC/MS. The p-values of these comparisons for individual proteins are then transformed into -log10 scale for volcano plot. Differential expression of each protein was normalized with Z-score transformation across all samples for heatmap visualization. z = (X – μ) / σ where z is the Z-score, X is the abundance of each protein detected by LC/MS, μ is the mean of the abundance for each protein across each time point and cell line, and σ is the standard deviation. Proteins that maximally expressed in a cell line at each time point are binned together and then sorted in descending order^43^ to visualize (seaborn package in python) the patterns of protein expression change across the time points for each of the cell lines. Enrichment of biological pathway was analyzed with Reactome Knowledgebase (https://reactome.org).

### Three-Dimensional Cell Culture

Single cell suspensions of 2000 cells of T-REx 293 cell lines stably integrated with Con1-SPOP, Con1-SPOP_mut_ and Con1_mut_-SPOP were seeded per 96-well of the ultra-low attachment plate (Corning). The plate is centrifuged clockwise at 300 x g for 5 min and anti-clockwise for 5 min to allow aggregation of the cells. The cells were incubated at 37°C, 5% CO_2_ and 90% relative humidity and form spheroids until the diameter of individual spheroids reaches about 300 μm. Fresh media with or without 100 ng/mL doxycycline were added to the spheroid to allow expression of the constructs.

For the CellTiter Glo 3D (Promega) assay, after 80 hr incubation, equal volumes of the assay reagent were added as per manufacturer’s instruction and incubated for 30 min at room temperature with shaking. Luminescence was recorded using the Tecan M1000 plate reader.

### Flow cytometric analysis

Cells were seeded in 6-well poly-D-lysine coated plates and allowed to incubate overnight. The media was removed and replaced with serum-free media, followed by Lipofectamine-Messenger MAX (ThermoFisher) mediated transfection of vhhGFP-SPOP and its control mRNAs. 24 hr after transfection, cells were trypsinized and resuspended in cold PBS containing 10% FBS. The cell suspension was passed through a 35 μm nylon mesh to dissociate aggregates before analysis on BD LSRFortessa™ X-20.

### Immunofluorescence imaging and confluency measurements

Cells were seeded in 96-well poly-D-lysine coated μCLEAR® plates (Greiner) and allowed to attach overnight. The next day, transfection and dox-induction were performed as described above. For immunostaining to detect γH2A.X and PCNA, cells were fixed in 4% formaldehyde in PBS for 15 min and blocked with 5% normal donkey serum and 0.3% Triton X-100 in PBS for 1 hr at room temperature. Rabbit anti-γH2A.X (Ser139) antibody (Cell Signaling Technology, #9718) and mouse anti-PCNA (Cell Signaling Technology, #2586) was diluted in 1% BSA with 0.3% Triton™ X-100 in PBS and incubated overnight at 4°C. The next day, donkey anti-rabbit Alexa Fluor 647 (ThermoFisher, A-31573) and donkey anti-rabbit Alexa Fluor 488 (ThermoFisher, A-21206) was added for 1 hr at room temperature. Nuclei were counterstained with Hoechst. Images of fixed cells were acquired at 37°C, 5% CO_2_ using the Opera Phenix™ High Content Confocal Screening System under the 20X water immersion lenses. For confluency measurements, the percentage confluency of cells were tracked continuously over 4 days using the IncuCyte® S3 Live-Cell Analysis System under the 10X imaging objective and analyzed using the IncuCyte® software.

### *In vivo* efficacy in T-REx 293 xenograft mouse smodel

The protocol and any amendment(s) or procedures involving the care and use of animals in this study were reviewed and approved by the Institutional Animal Care and Use Committee (IACUC) of concerned institutions prior to conduct. During the study, the care and use of mouse were conducted in accordance with the regulations of the Association for Assessment and Accreditation of Laboratory Animal Care (AAALAC). This study was performed at HD Biosciences, San Diego. 7-8-week old female NIH-III nude mice (Charles River) weighing approximately 18-22 g were inoculated subcutaneously into the right lower flank in 0.1 mL of serum-free MEM medium mixed with basement membrane Matrigel (in a 1:1 ratio) for tumor development. The total number of cells implanted for each cell line (Parental T-rex-293, Con1-SPOP, Con1_mut_-SPOP and Con1-SPOP_mut_) in the right side was 10 x 10^6^ cells per mouse. Tumor starting size and range for efficacy cohort and PD was approximately 50 mm^3^. Doxycycline treatment (Doxycycline Hyclate USP 625mg/kg chow, Envigo) was started when at least 11 mice per cell line reached a mean tumor size of approximately 50 mm^3^ (40-60 mm^3^ range). Amount of food weight consumed was measured weekly. Tumors were measured twice a week in two dimensions using a caliper, and the volume was expressed in mm^3^ using the formula: V = 0.5 (a x b^2^) where a and b are the long and the short diameters of the tumor, respectively. Body weights were taken twice/week. Mice with complete regression (no measurable tumor) were followed until the end of study date. PD tumor samples were collected 3 days after Dox diet (G1a, G2a, G3a and G4a). Tumors will be collected and cut in half and were homogenized for Western blot. Western blot analysis was performed with FLAG (Sigma-Aldrich F1804-1MG), PCNA (Cell Signaling Technology #13110), Cleaved Caspase 3 (Cell Signaling Technology #9664) and GAPDH (Cell Signaling Technology #5174) antibodies.

### *In vitro* transcription of mRNA and LNP preparation

mRNAs were *in vitro* transcribed using the mMESSAGE mMACHINE® T7 Ultra transcription kit (Ambion, AMB13455) or synthesized at TriLink Biotechnologies, capped with CleanCap® and modified with 100% pseudouridine. cDNA encoding biodegrader was cloned into pEF6 with NheI and NotI sites. Linearized plasmid DNA containing the target gene downstream of a T7 RNA polymerase promoter was used as the template and synthesis reactions were performed according to the manufacturer’s protocol. mRNAs were subsequently purified by the RNeasy Mini Kit (Qiagen, #74104) and concentration quantified on the NanoDrop spectrophotometer by Abs260. RNA quality was checked with 1% agarose gel. For LNP preparation, lipids were dissolved in ethanol and mixed in weight ratio of 1:16 for mRNA:lipid. The aqueous and ethanol solutions were mixed in a 3:1 volume ratio using a microfluidic device NanoAssemblr (Prevision NanoSystems). Formulated LNP was dialyzed overnight with Tris buffer. The encapsulation efficiency of mRNA was determined using the RiboGreen assay. Pseudouridine modified Con1-SPOP and Con1_mut_-SPOP mRNA were encapsulated. The final ratio of lipids by percentage was 58:30:10:2 (ionizable cationic lipid:Cholesterol:Phospholipid:PEG-lipid).

### Transfection of mRNA, siRNA and western blot analysis

Messenger RNA transfection follows Lipofectamine™ MessengerMAX™ (ThermoFisher, #LMRNA001) transfection protocols. Briefly, Con1-SPOP and Con1_mut_-SPOP mRNA (TriLink) were diluted with Opti-MEM™ Medium follow by addition of diluted MessengerMAX reagent at 1:1 ratio. The mRNA-lipofectamine complex was added to cells in Opti-MEM™ Medium and allow to incubate for 4 hr, follow by addition of complete culture media. The PCNA siRNA was acquired from Dharmacon through the ON-TARGETplus siRNA platform. siRNA transfection follows DharmaFECT™ transfection protocol. Briefly, PCNA siRNA, siGAPDH and siScramble controls were diluted with the appropriate DharmaFECT transfection reagent with serum free medium according to manufacturer’s instructions and allow to incubation at room temperature for 20 mins. The culture media from the wells of 12-well plate was removed and 1000 μl of the respective transfection medium was added to each well with siRNA at 25 nM final concentration. The cells were incubated for 72-96 hr for protein analysis. Thawed LNPs or LNPs loaded with biodegrader or non-binding control mRNA were diluted to appropriate concentrations with complete media and allow to incubate at room temperature for 10 mins and added directly to cells incubated in complete media. Cells were lysed in ice-cold RIPA lysis buffer (ThermoFisher, #89900) supplemented with cOmplete™ EDTA-free protease inhibitor cocktail (Roche, # 11836170001) and Phosphatase inhibitor (Roche, #4906837001) for 30 min with intermittent vortexing. Lysates were centrifuged at 18,000 g, 4°C for 15 min and supernatants were snap frozen in liquid nitrogen. Protein concentration was determined using the BCA protein assay kit (ThermoFisher, #23225). 20 to 30 μg of protein extract was separated on 4-12% Bis-Tris plus gels, transferred onto nitrocellulose membranes using the Trans-Blot® Turbo™ semi-dry system (Biorad), and blocked for 1 hr at room temperature with tris-buffered saline (TBS) Odyssey blocking buffer (Li-Cor). Blots were probed with the appropriate primary antibodies overnight at 4°C in Odyssey blocking buffer supplemented with 0.1% Tween-20, followed by the secondary antibodies IRDye® 680RD donkey anti-mouse IgG and IRDye® 800CW donkey anti-rabbit IgG (Li-Cor) for 1 hour at room temperature. Fluorescent signals were imaged and quantified using Odyssey® CLx. Primary antibodies used were: DNA damage antibody sampler kit (Cell Signaling Technology, #9947), PLK1 (Cell Signaling Technology, #4513), WEE1 (Cell Signaling Technology, #13084), Hsp90 (BD Transduction Laboratories, #610419), FLAG-tag (Cell Signaling Technology, #8146 and #14793), PCNA (Cell Signaling Technology, #2586 and #13110).

## ACKNOWLEDGEMENTS

We thank Eric Gifford from MRL IT Singapore and all members of the Quantitative Biosciences team, U-ming Lim and Aaron Fernandis from Target & pathway biology for helpful discussions and comments on the experimental design and manuscript. We also thank Byung Lee and the team at HD Biosciences for their contribution to the *in vivo* study. The authors acknowledge support from the MRL Postdoctoral Research Program.

## SUPPLEMENTARY INFORMATION

**Figure S1.**
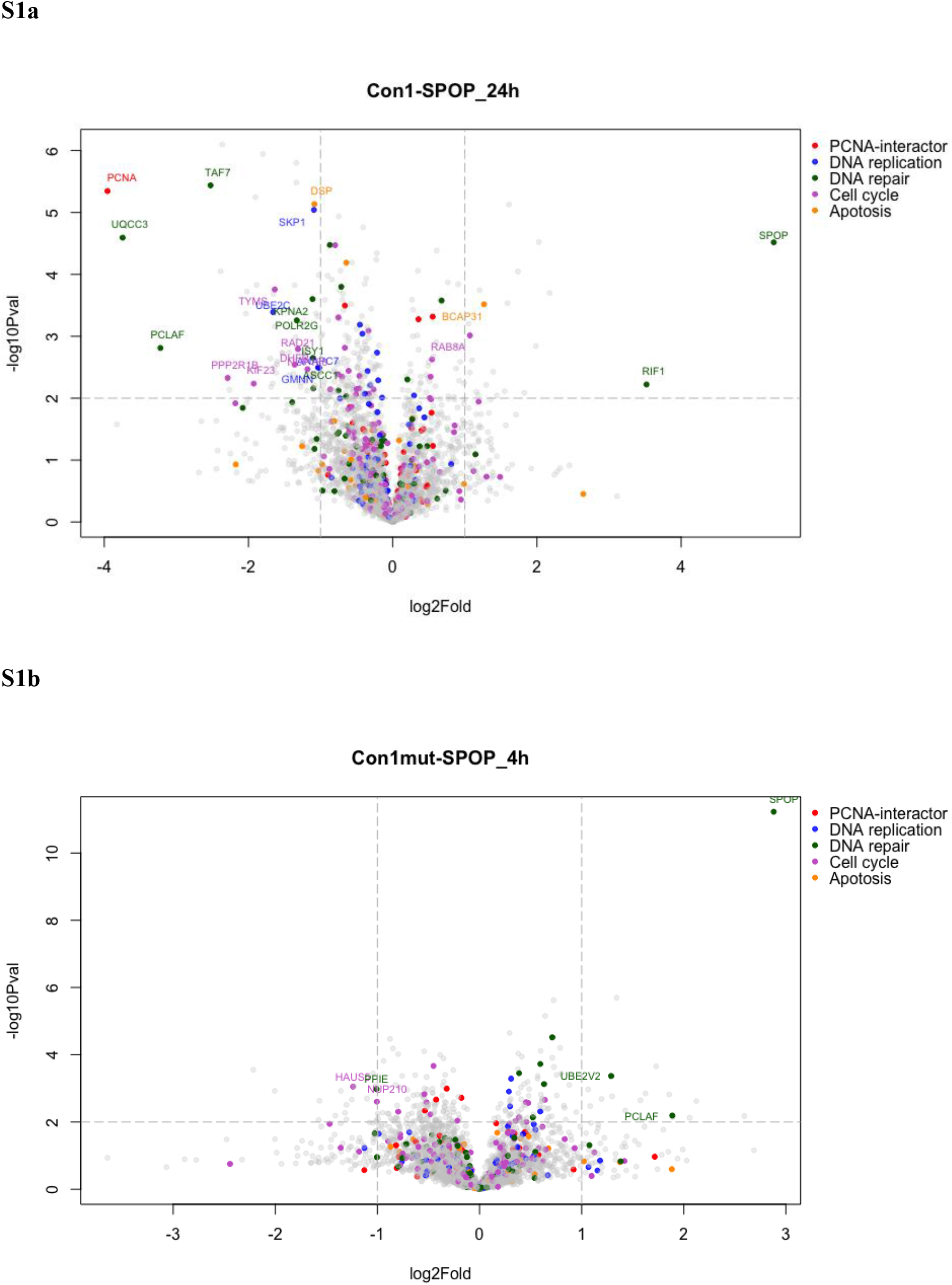
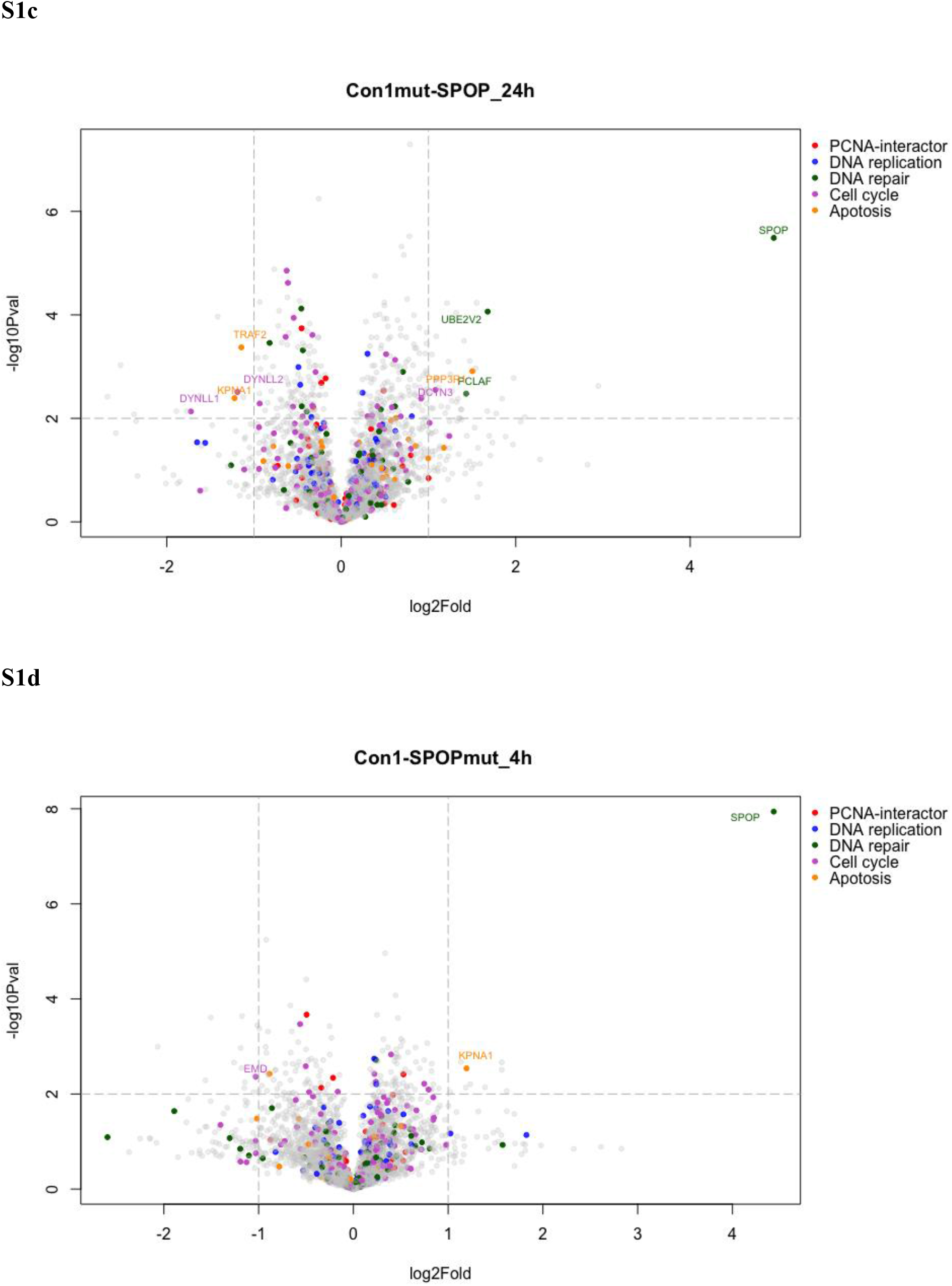
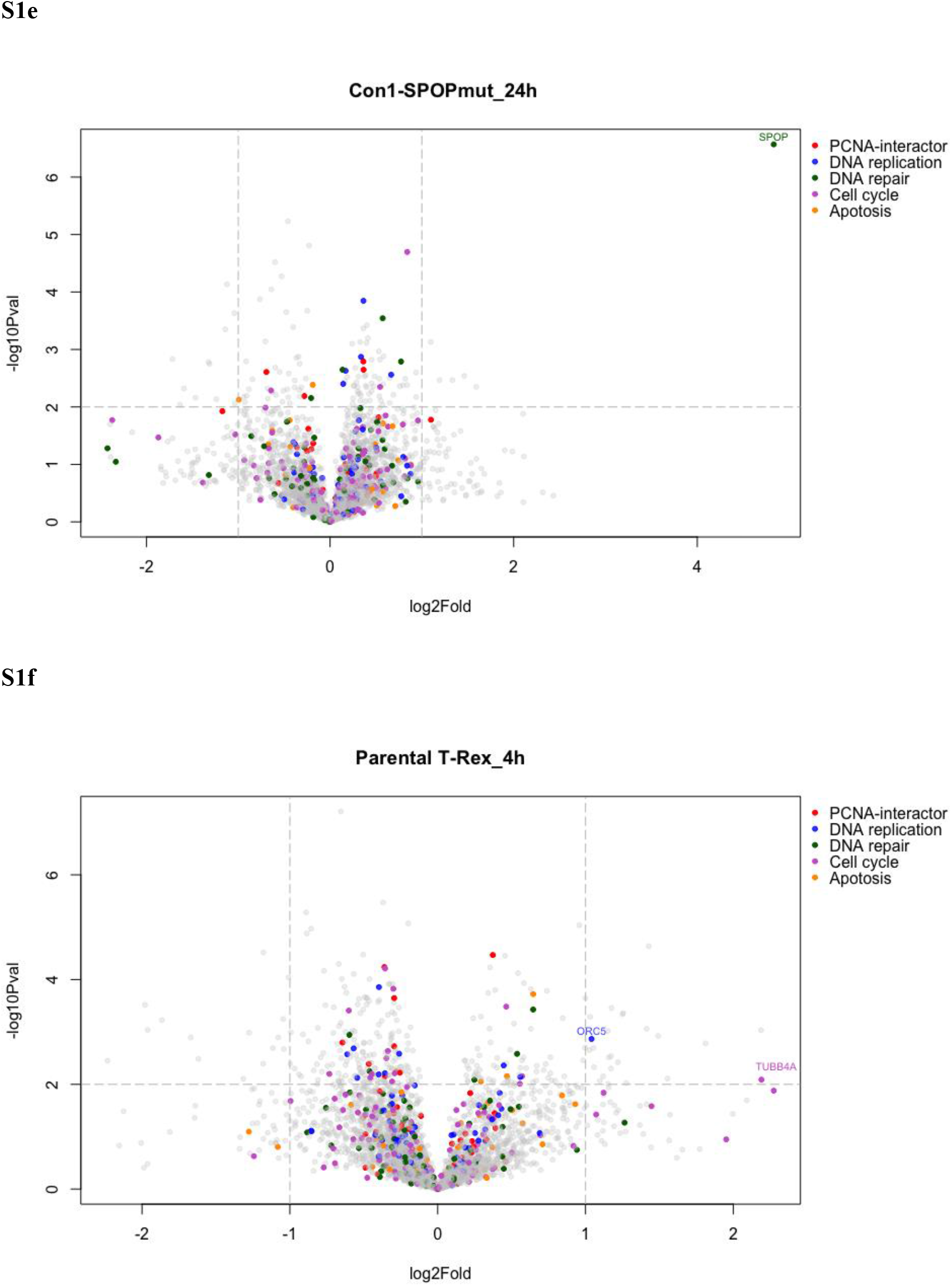
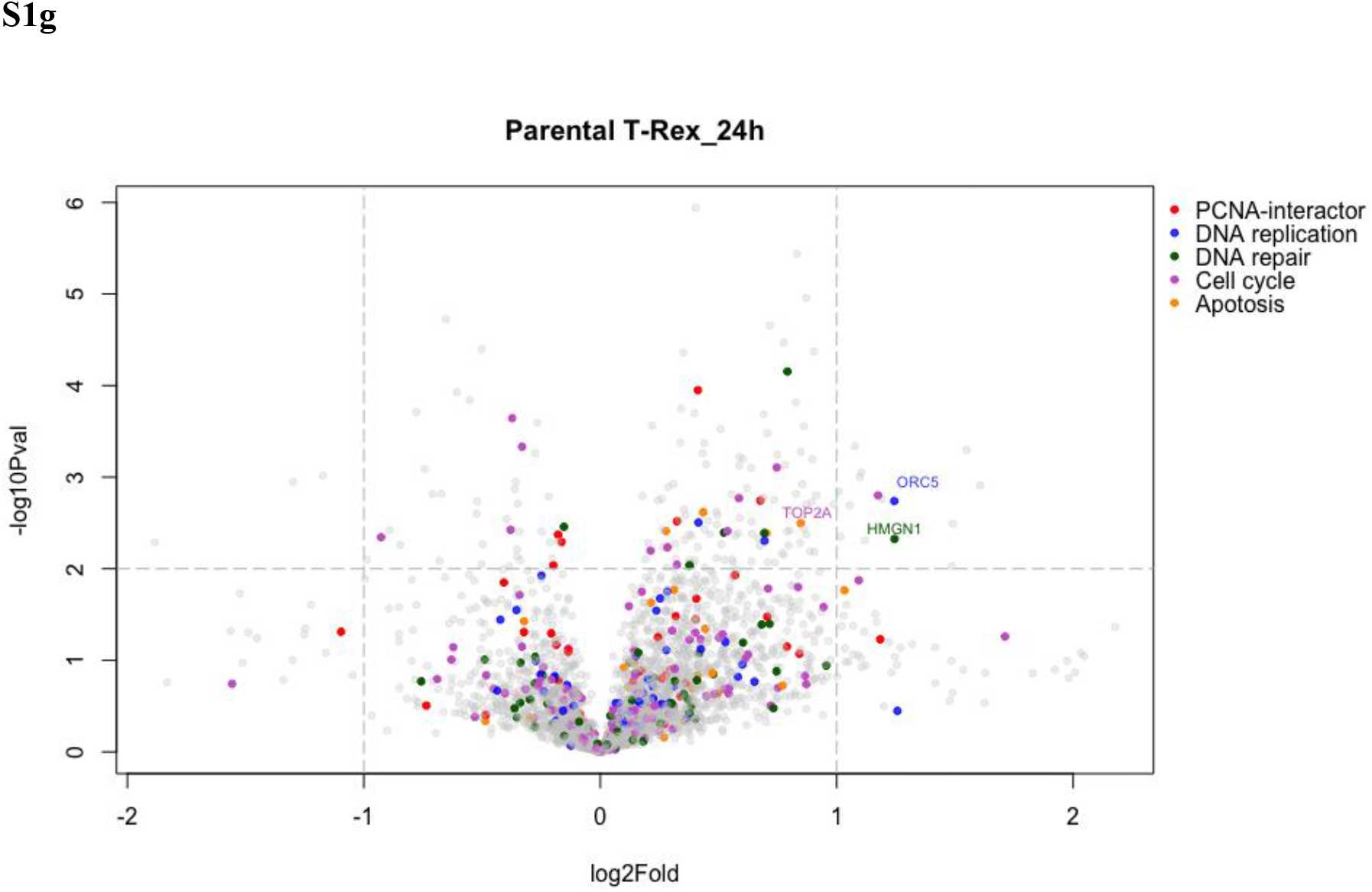
Volcano plots of protein differentials of T-REx-293 cell lines induced with 100ng/ml of doxycycline as compared to no Dox control. (a) Con1-SPOP at 24 hr, (b) Con1-SPOP_mut_ at 4 hr, (c) Con1-SPOP_mut_ at 24 hr, (d) Con1_mut_-SPOP at 4 hr, (e) Con1_mut_-SPOP at 24 hr, (f) Parental T-REx-293 at 4 hr and (g) Parental T-REx-293 at 24 hr. Proteomic analysis is based on 3 independent replicate experiments.

**Figure S2.**
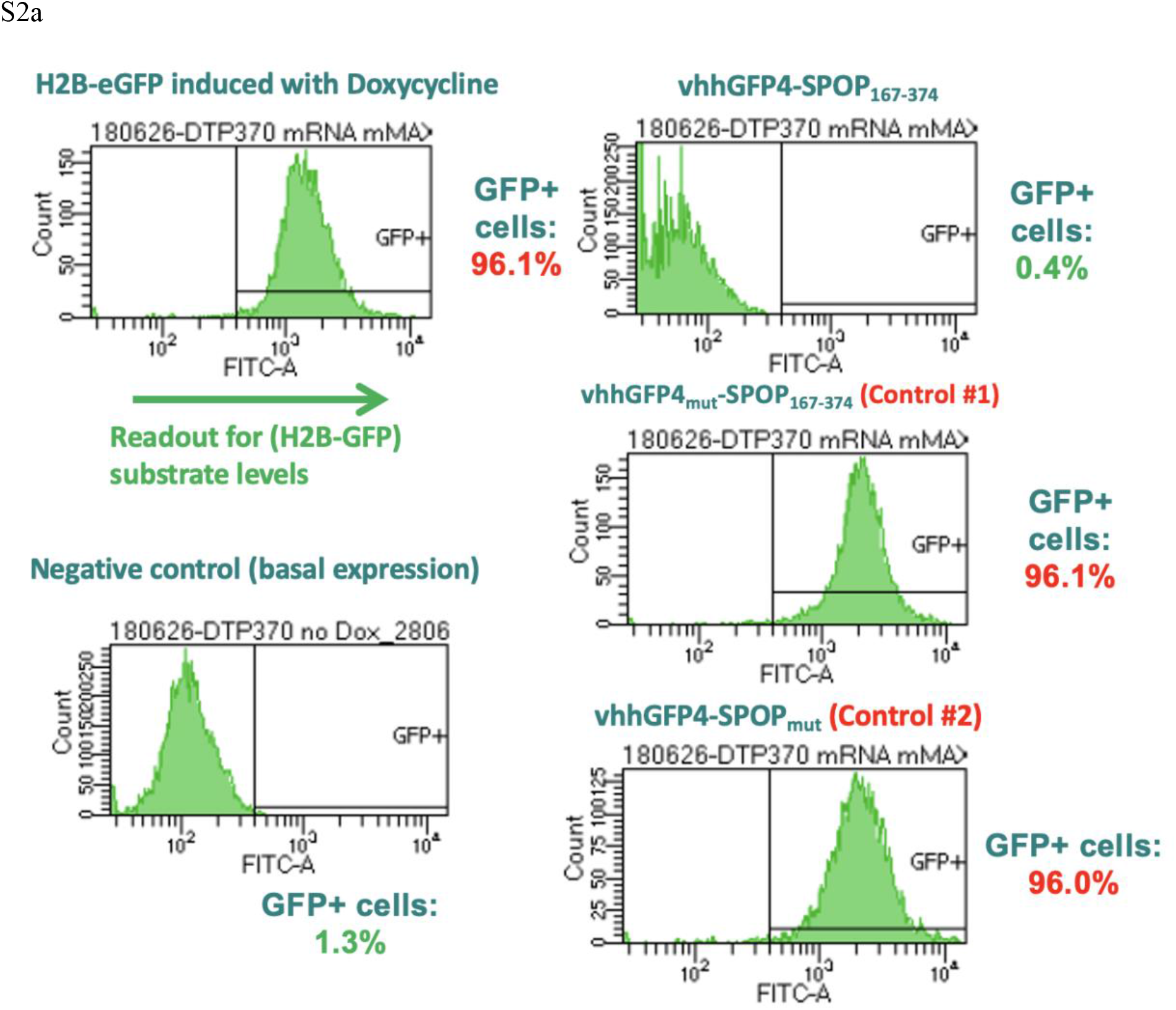
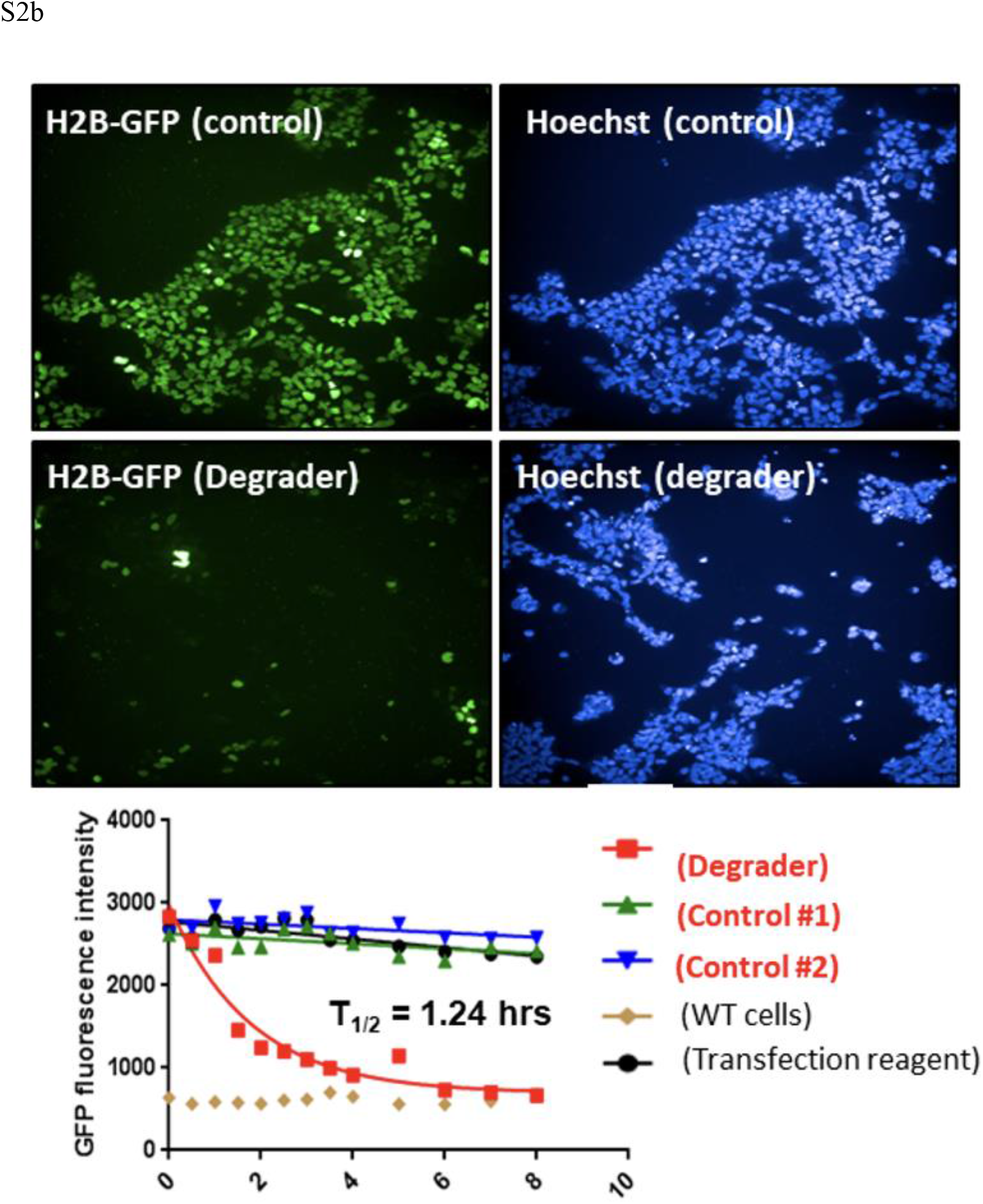
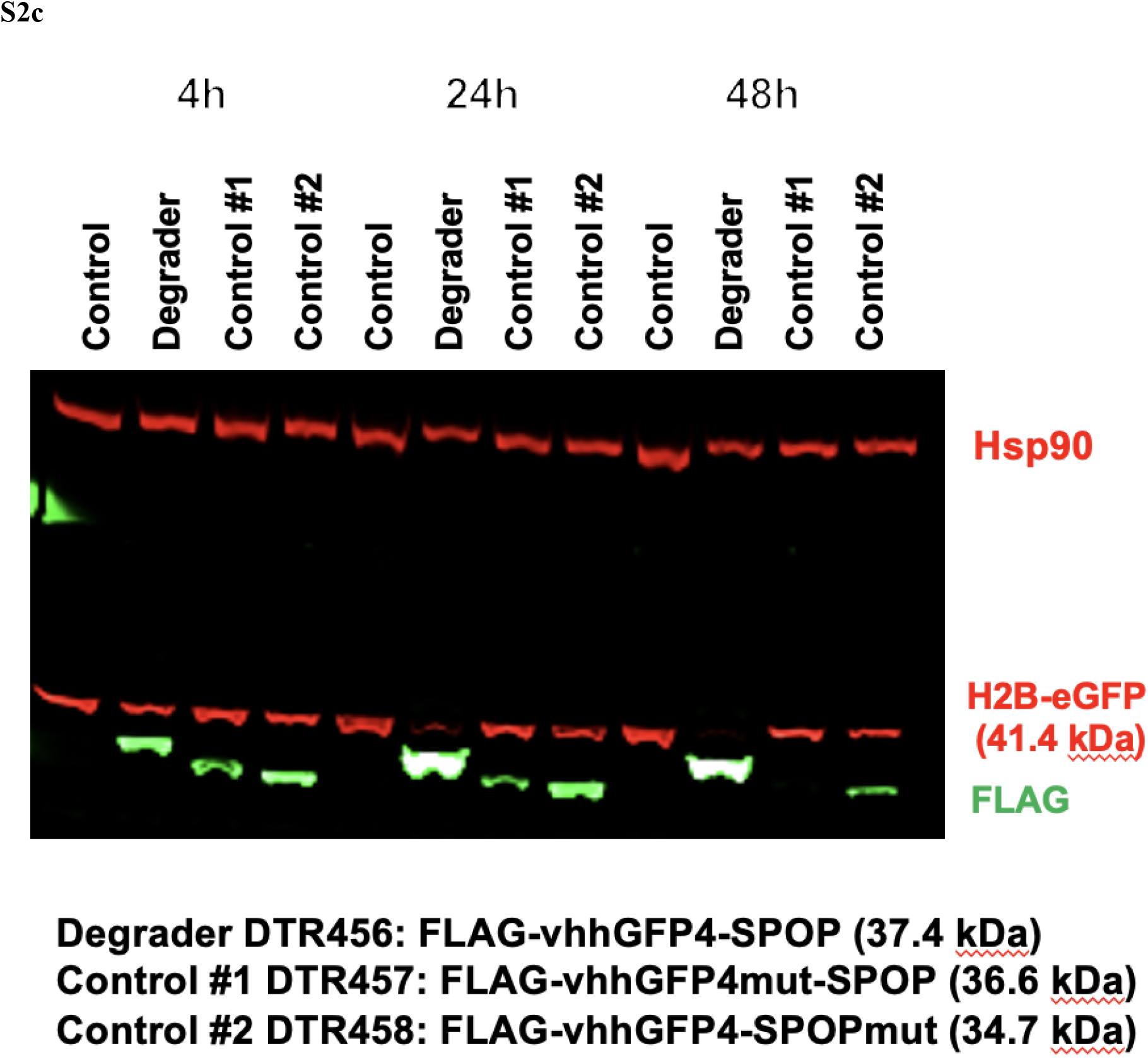
Degradation of H2B-GFP fusion protein by GFP biodegrader. (a) Flow cytometry analysis of GFP fluorescence of Dox-inducible H2B-GFP stable cell line transfected with vhhGFP-SPOP and its control mRNAs, demonstrating high degradation efficiency of H2B-GFP. (b) Confocal imaging of Dox-inducible H2B-GFP stable cell line transfected with vhhGFP-SPOP mRNA and measurement of the half-life of H2B-GFP fluorescence. c) Western blot analysis of H2B-GFP degradation at 4h, 24h and 48h.

**Figure S3.**
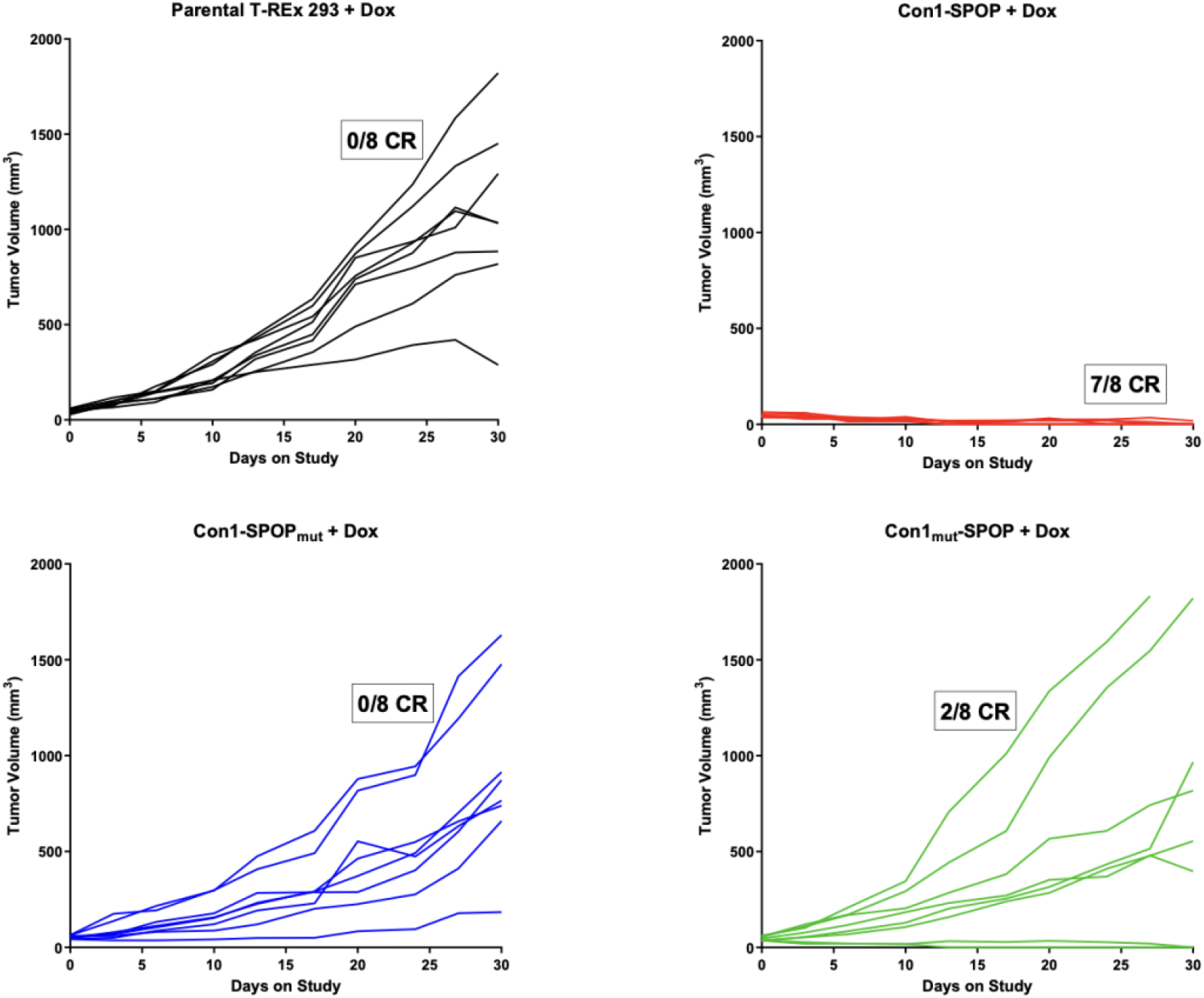
Individual tumour volumes of Dox-induced Con1-SPOP and its respective controls in the T-REx 293 xenograft model. Tumor volumes were measured twice a week after Dox-treatment until Day 30 for each group (*n* = 8): parental T-REx 293 or T-REx 293 cells expressing Con1-SPOP, Con1-SPOP_mut_ or Con1_mut_-SPOP. For each group, number of mice that showed complete tumor regression (CR) is indicated.

**Figure S4.**
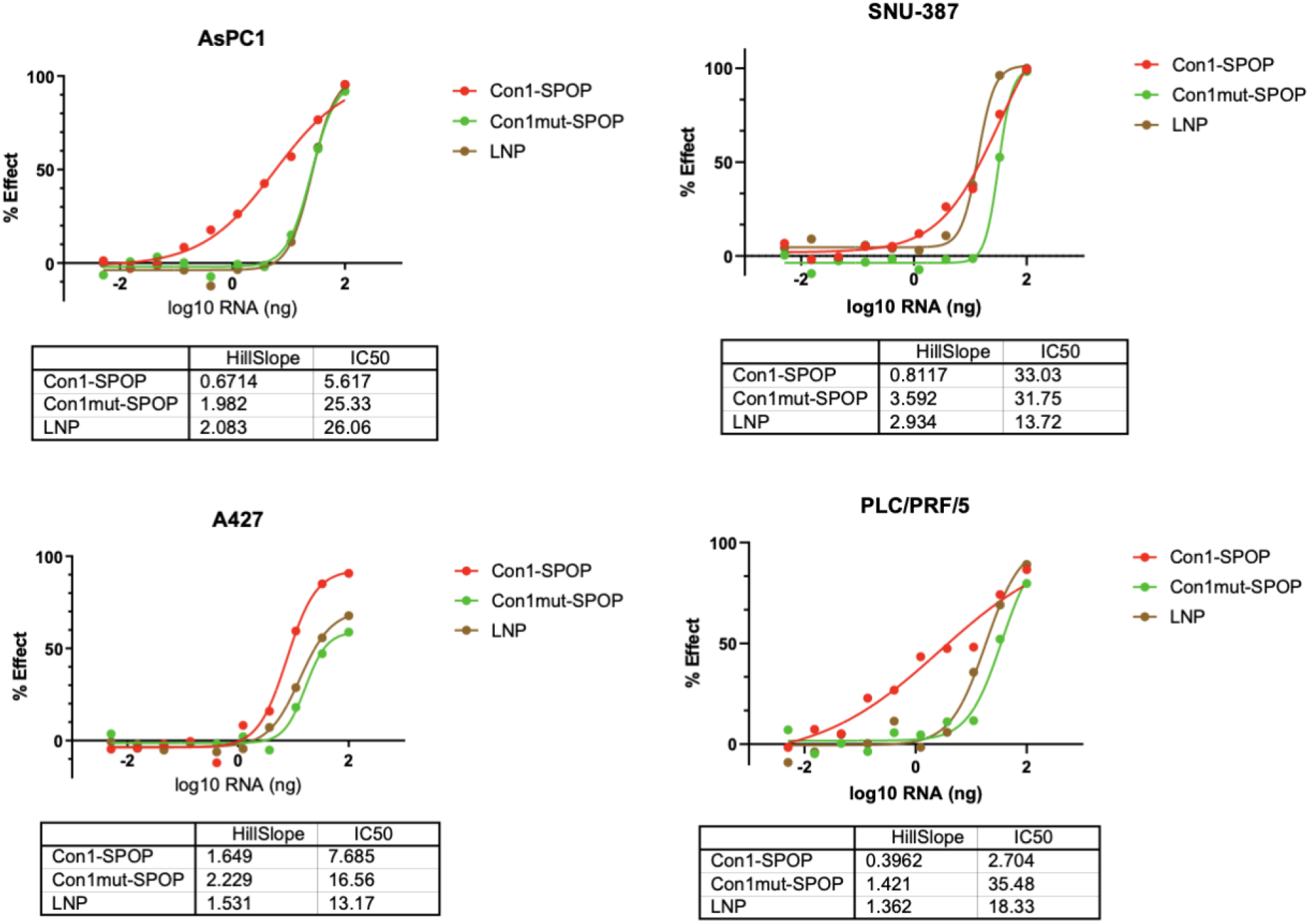
Quantification of cytotoxicity of A427, AsPC-1, PLC/PRF/5, SNU-387 with LNP-Con1-SPOP (or its control). Cell viability determination using CellTiter-Glo assay performed for respective cell line at 72 hr post treatment with a dose titration of empty LNPs or LNPs loaded with biodegrader or non-binding control mRNAs. Data was normalized to untreated controls in respective cell lines.

